# Subnuclear cofactor partitioning underlies auxin-dependent transcriptional regulation

**DOI:** 10.64898/2026.04.29.721720

**Authors:** Jorge Hernandez-Garcia, Melissa Dipp-Alvarez, Martijn de Roij, Maria Guillem-Bernal, Mark Roosjen, Zeynep Baltaci, Harrison W. Smith, Danilo Pereira, Jan Willem Borst, Ales S. Holehouse, Hyon O. Lee, Dolf Weijers

## Abstract

Cellular and organismal function relies on the precise activation and repression of gene expression by DNA-binding transcription factors (TFs). Many TFs occur in large gene families, and a key question in biology is how divergent functions emerge in TF families during evolution. Here we discover the biochemical mechanism for transcriptional activation by the *Marchantia polymorpha* AUXIN RESPONSE FACTOR1 (MpARF1) TF, which relies on direct recruitment of the Mediator complex into subnuclear MpARF1 clusters. We find that the Mediator recruitment region was the evolutionary innovation that converted ARF repressors into activators, switching binding specificity from co-repressor to co-activator. We demonstrate that this evolutionary innovation can be recreated, thereby revealing a deeply conserved mechanism based on competition between ARF clusters at the heart of the transcriptional auxin response.

## Introduction

Transcriptional regulation is a highly coordinated process that involves both activation and repression to maintain precise control over gene expression. Transcriptional activation relies on transcription factor (TF) binding to specific DNA motifs in gene regulatory regions, and the subsequent recruitment of co-activators and RNA polymerase II, to initiate transcription (*1–4*). Conversely, transcriptional repression requires TFs to recruit co-repressors that locally modulate chromatin to actively prevent transcription (*2, 5–9*). The dynamic interplay between activating and repressive mechanisms ensures that genes are expressed only in the appropriate cellular context, developmental stage, or in response to environmental cues, a key requirement for organismal function (*10–12*).

An excellent example of the coordination between transcriptional activation and repression is the regulation of auxin-dependent gene expression. Auxin is a key signalling molecule that controls crucial aspects of growth and development in land plants, in large part through a nuclear auxin signalling pathway that controls gene expression (*13*). Transcriptional regulation is orchestrated by the AUXIN RESPONSE FACTOR (ARF) TFs, which function as either transcriptional activators (A-class) or repressors (B-class)(*14, 15*). Auxin-mediated control of transcription depends on the antagonism between these ARFs, since both compete for the same DNA binding sites in the genome, and their stoichiometry determines the outcome of the auxin signal (*16–18*).

Interestingly, these antagonistic TFs emerged from a gene duplication event of a single ancestral copy encoding a repressor ARF (A/B-ARF) (*19, 20*). A key question is how the transcriptional activation function, which defines the heart of auxin response, evolved. This question is difficult to address given that no general biochemical mechanism for ARF-triggered transcriptional activation has yet been reported. Here, by dissecting the requirements and molecular mechanism underlying transcriptional activation by the single A-class ARF (MpARF1) in *Marchantia polymorpha* (henceforth “Marchantia), we address how a transcriptional activator evolved from repressors and gave rise to the core of the transcriptional auxin response system.

## Results

### Identification of a minimal, essential ARF transcriptional activation domain

The acquisition of transcriptional activation functions occurred in the land plant-specific A-class ARF clade (*20*), which has substantially expanded and diversified in flowering plants (*19*). Previous studies in *Arabidopsis thaliana* have shown different partner proteins to mediate the activity of different A-ARFs subclades. These include Mediator subunits, SWI/SNF2 chromatin remodelers, and histone methyltransferase ATXR2 (*21–23*). To identify core, ancestral mechanisms preceding paralog functional divergence, we previously studied the liverwort *Marchantia polymorpha* that carries a minimal auxin transcriptional system with single A-activation mechanism (*14*). Existing prediction tools for plant activation domains failed to highlight a robust candidate region (see methods, fig. S1A) (*24*). Using a yeast-one hybrid-based transcriptional assay, we experimentally identified an irreducible 106 amino acid region (F2) within the intrinsically disordered Middle Region (MR) that is both necessary and sufficient for transcriptional activation in the assay (**Fig. 1A**; fig. S1B). Analysis of the F2 region protein sequence revealed signatures of a single predicted prion-like domain (PrD, LLR = 42.03), fully contained within the region, followed by a stretch enriched in proline, serine and leucine residues (binomial enrichment *versus* MR, p = 2.5 × 10^-4^, Table S1), embedded within the MR (**Fig. 1B**). Neither the B-class MpARF2, nor the C-class MpARF3 presented PrD signatures in their sequence (fig. S2), indicating that PrD features are specific to MpARF1, and B-class ARFs (*16*). Here, we use its A-class, MpARF1, to dissect the elusive ARF transcriptional and thus correlate with transcriptional activity. Indeed, while the MR of MpARF1 activated transcription in plant cells, a version without F2 retained only residual activity (**Fig. 1C**).

**Fig. 1.**
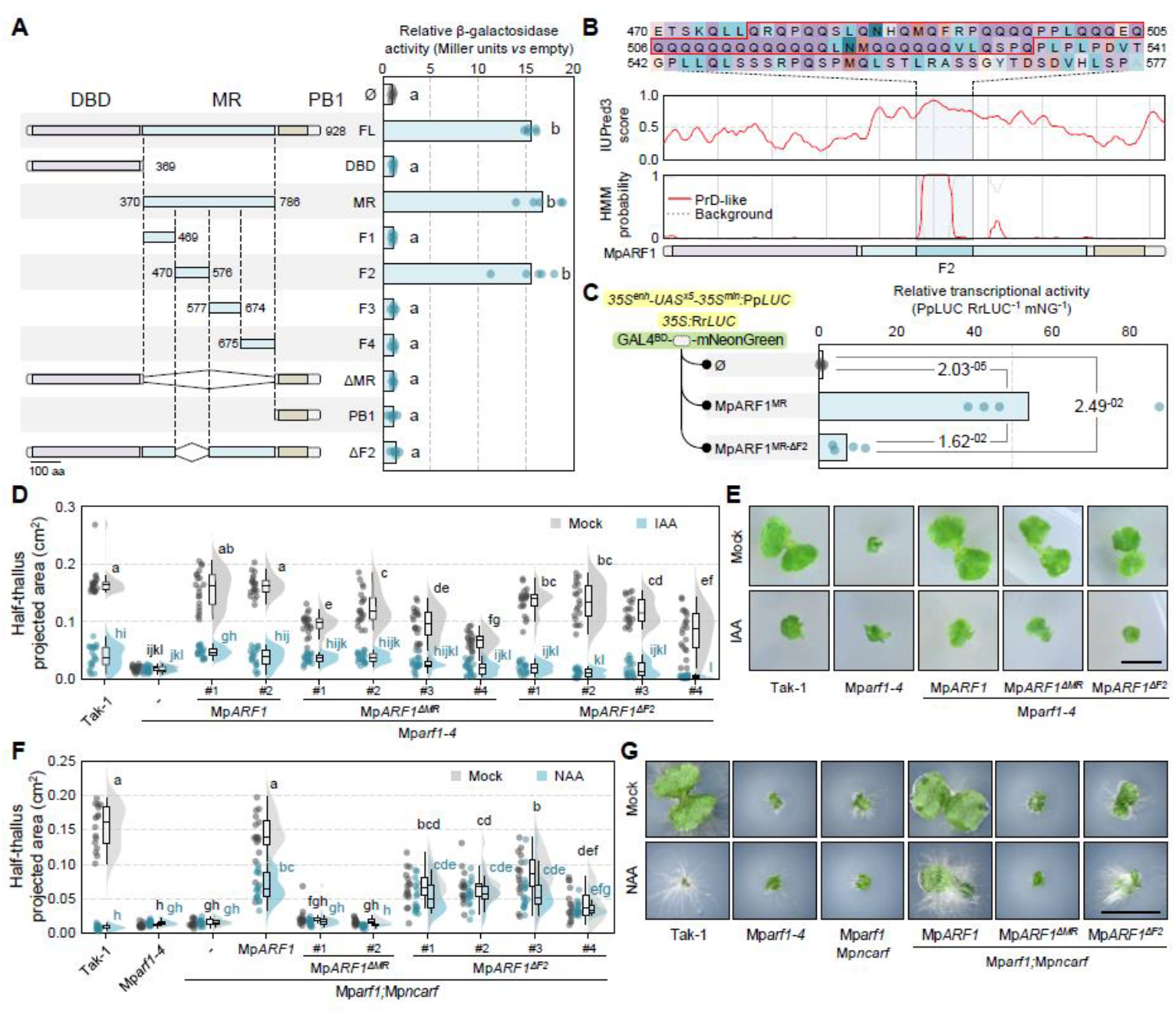
Identification of a minimal transcriptional activation domain in MpARF1. **(A)** Transcriptional activation as a measure of β-galactosidase activity (Miller units) of MpARF1 and fragments thereof in a yeast one-hybrid assay. Amino acid positions are indicated. DBD, DNA-binding Domain; MR, Middle Region; PB1, Phox-Bem1. **(B)** Prediction of disorder (IUPred3) and Prion-like domains (HMM) in MpARF1. The sequence of the F2 region is indicated on top. **(C)** Transcriptional activation by MpARF1-MR and an F2 deletion in an Arabidopsis protoplast-based assay. **(D,E)** Complementation of Mp*arf1-4* mutant by MpARF1, as well as MpARF1 versions with MR or F2 region deleted. Plants were grown on 3 µM auxin (IAA) or mock media (DMSO) and growth **(E)** was quantified as half-thallus projected area. **(F,G)** Complementation of Mp*arf1;*Mp*ncarf* double mutant by MpARF1, as well as MpARF1 versions with MR or F2 region deleted. Plants were grown on 3 µM synthetic auxin (NAA) or mock media (DMSO) and growth **(G)** was quantified as half-thallus projected area. Scale bars are 5 mm in **(E)** and **(G)**. In **(A)**, statistical groups are determined by Dunn’s test after Kruskal–Wallis H test (*p* < 0.05). In **(C)**, statistical differences are determined after log-transformation of data followed by pairwise Welch’s t-test with Holm correction for multiple testing. In **(D,F)** boxplots in rainclouds indicate the following parameters: centrum, median; upper bound, first quartile; lower bound, third quartile; whiskers maximum and minimum refer to highest and lowest values, respectively, within 1.5*inter-quartile range (IQR). Statistical groups are determined by Tukey’s Post-Hoc test (*p* < 0.05) following one-way ANOVA.

To explore the biological relevance of this fragment, we performed genetic complementation assays of the Mp*ARF1* knock-out mutant, Mp*arf1-4 (25*) using native regulatory elements (**Fig. 1D,E**). This mutant is insensitive to auxin and displays stunted growth compared to wild-type plants. We first introduced an Mp*ARF1* copy with its entire MR deleted (Mp*ARF1*^*ΔMR*^). This led to partial complementation of its phenotypes (**Fig. 1D,E**). Introducing a Mp*ARF1* copy with the F2 fragment deleted (Mp*ARF1*^*ΔF2*^) resulted in an intermediate partial complementation (**Fig. 1D,E**), showing that F2 is relevant for *in vivo* function. As expected, Mp*ARF1*^*ΔF2*^ lines showed impaired expression of known MpARF1-regulated genes (fig. S3A).

The partial complementation is remarkable, since no elements for transcriptional function were found outside the MR (**Fig. 1A**), and transcriptional activation of its targets would be expected to be an essential MpARF1 function. We reasoned that back-up transcriptional co-activation may be provided by the non-canonical ARF (ncARF; also referred to as D-class ARFs) proteins representing an A-class-derived clade, lacking a DNA-binding domain (*19, 26*). We found that the single Marchantia ncARF (MpncARF) could indeed activate transcription in yeast, and interact through their PB1 with MpARF1 (fig. S3B). This suggests a model where ARF1 DBD binds DNA, and ncARF associates through PB1 interactions to boost transcriptional activity, and the ncARF F2 domain would compensate for the loss of the MpARF1 F2 domain. Indeed, deleting the ARF1 DNA-binding domain renders the protein entirely inactive (fig. S4). Therefore, we generated Mp*arf1-4;*Mp*ncarf* double mutants, which phenocopied Mp*arf1* (**Fig. 1F,G**). A copy of ARF1 lacking

The partial complementation is remarkable, since no elements for transcriptional function were found outside the the MR ((Mp*ARF1*^*ΔMR*^) completely failed to complement the Mp*arf1-4;*Mp*ncarf* double mutant (**Fig. 1F,G**), supporting the idea that ncARF acts to supplement transcriptional activity of ARF1. Deleting only F2 led to partial complementation, but eliminated the response to auxin (**Fig. 1F,G**). Hence, the transcriptional function endowed by F2 is essential for auxin-dependent ARF1 activation, yet there may exist an activation- and F2-independent function in the MR.

### MpARF1 directly recruits the Mediator complex to promote transcriptional activation

A key question is through what mechanisms ARFs promote transcription. We capitalized on the identification of F2 as a necessary and sufficient region for activation to explore which proteins may mediate F2 function. We generated transgenic lines carrying either MpARF1 or MpARF1^ΔF2^ tagged with TurboID for biotin-based proximity labelling. These fusion proteins localized to the nucleus and recapitulated the phenotypes expected from functional proteins (figS5). We performed proximity labelling, treating with auxin to enrich activation-related complexes. In addition to knownauxin-signaling components, the MpARF1-TurboID proxitome was enriched in several Mediator tail subunits (MED3,5,16,23) (**Fig. 2A**, Data S1), known to mediate TF-specific recruitment of the entire Mediator complex (*27, 28*). In addition, we found enrichment in MED14, suggesting the presence of fully assembled complexes (*29*). In contrast, the proxitome of TurboID-tagged MpARF1^ΔF2^ showed low-to-no enrichment in these Mediator subunits, with the sole exception of MED8, which appeared less enriched in MpARF1 (**Fig. 2A,B**). Notably, several of the subunits (MED5,14,28) were absent from the MpARF1^ΔF2^ proxitome. Likewise, subunits from other transcriptional complexes, such as the SWI/SNF2 (Swp73 and ARP4/7) or SAGA/NuA4 (MpTRA1), and several TFs associated with MpARF1 in an

**Fig. 2.**
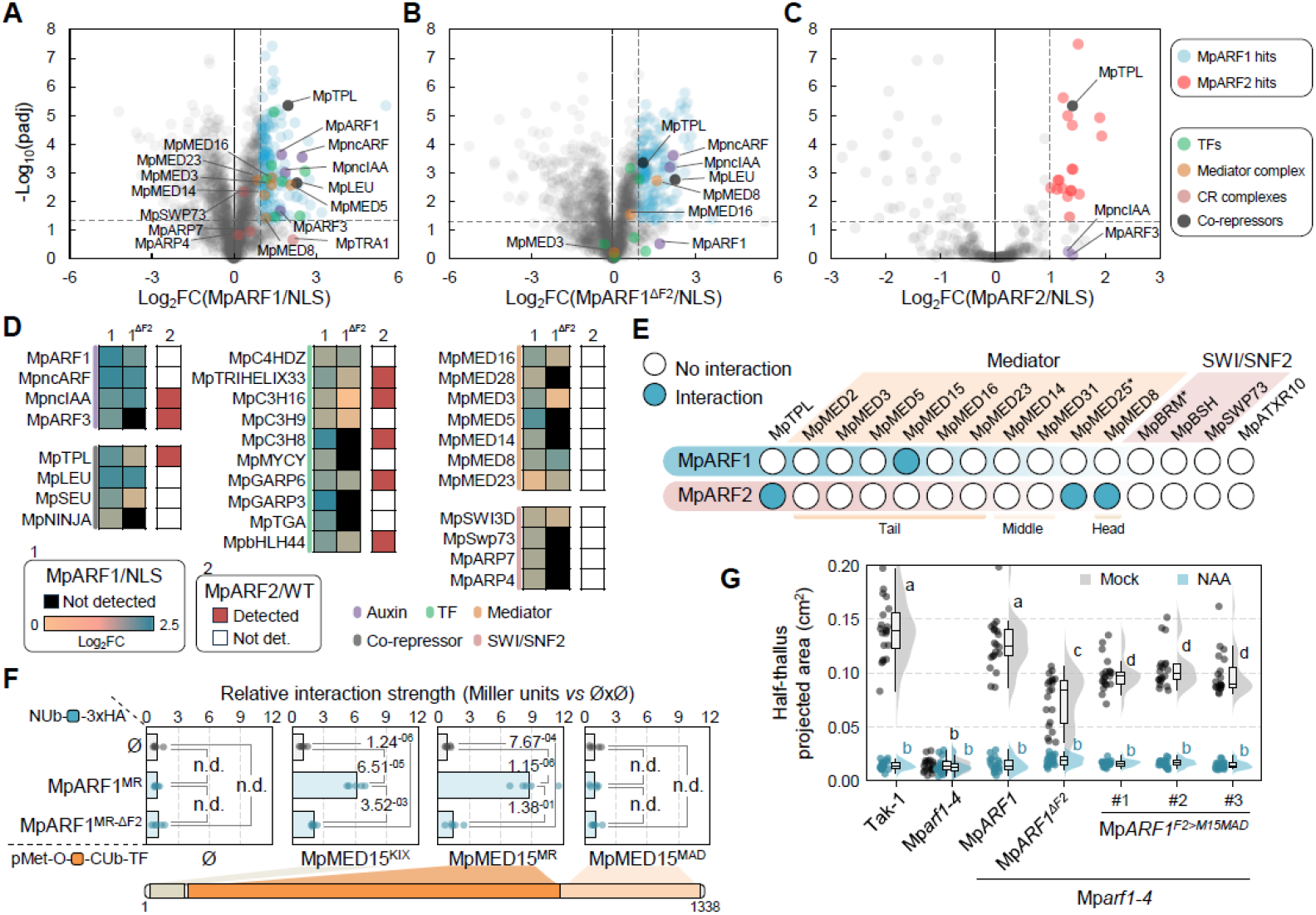
MpARF1 activates transcription by directly recruiting the Mediator complex. **(A-C)** Volcano plots of proximity ligation experiments on nuclei isolated from MpARF1-TurboID **(A)**, MpARF1ΔF2-TurboID **(B)** and MpARF2-TurboID (**C)** plants. Fold-enrichment relative to NLS-TurboID (Log2FC) is plotted against p-value across four technical replicates. Relevant proteins (legend) are indicated in the plots. **(D)** Enrichment of auxin response components, co-repressors, transcription factors, Mediator subunits, and chromatin remodelers in each TurboID experiment (1-MpARF1; 2-MpARF1ΔF2; 2-MpARF2), either as Log2FC or as absence/presence. **(E)** Summary of yeast cytosolic split-ubiquitin interaction assays between MpARF1 or MpARF2 and Mediator subunits, SWI/SNF2 subunits, and MpATXR10 (see **Fig. S6**). **(F)** Yeast cytosolic split-ubiquitin interaction assays between MpARF1MR or MpARF1MR-ΔF2 and MpMED15 domains. **(G)** Quantification of half-thallus projected area in Mparf1-4 mutant complemented with MpARF1, MpARF1ΔF2, as well as with MpARF1 where F2 is replaced by the MpMED15-MAD domain, grown on NAA and mock media. In **(E)**, positive interactions are determined by statistical difference against the empty prey in each case as determined by Dunn’s test after Kruskal–Wallis H test (p < 0.01). In **(F)**, statistical differences are determined after log-transformation of data followed by pairwise Welch’s t-test with Holm correction for multiple testing (p < 0.01). In **(G)**, boxplots in rainclouds indicate the following parameters: centrum, median; upper bound, first quartile; lower bound, third quartile; whiskers maximum and minimum refer to highest and lowest values, respectively, within 1.5*inter-quartile range (IQR). Statistical groups are determined by Tukey’s Post-Hoc test (p < 0.05) following one-way ANOVA.

F2-dependent manner (**Fig. 2A,B**). In contrast, proximity labelling using the repressive, B-class MpARF2-TurboID showed no Mediator or co-activator subunits, but verystrong enrichment of the TOPLESS co-repressor MpTPL (**Fig. 2C**, Data S1). We next explored possible direct interactions between MpARF1 and Mediator subunits using a yeast cytosolic split-ubiquitin assay. We first detected all the orthologs of the Mediator complex subunits encoded in Marchantia using a previously developed tool (See methods, Data S2) (*30*), and included all tail module subunits (MED2,3,5,15,16,23) and additional subunits from other modules (MED8,14,25,28,31) and previously proposed ARF-interacting subunits (*22*). In addition, we included additional SWI/SNF2 subunit orthologs previously reported as ARF partners (*31*) and the single Marchantia ATXR2 ortholog. We confirmed a direct interaction of MpARF1 with the Mediator tail subunit MED15 (**Fig. 2D**, fig S6). In contrast, MpARF2 was able to bind to MED8 and MED25 (**Fig. 2D**), previously shown to interact with A-ARFs (*21*), but known to play a dual role in transcription through their interaction with the co-repressor TPL (*32*). Further dissection of MpARF1 and MED15 interaction showed that F2 drives MpARF1 interaction with the disordered middle region of MED15, and to a lesser extent with the globular KIX domain (**Fig. 2E**).

We next tested whether the recruitment of the Mediator complex by MpARF1 contributes to biological function. To this end, we engineered a version of MpARF1 in which the F2 region was replaced by the Mediator Associated domain (MAD) of MpMED15. This domain is responsible for recruiting the rest of the Mediator complex (*33*), and should thus bypass the need for F2-MED15 interaction. Indeed, the Mp*arf1-4* mutant phenotype was partially rescued by this chimaera (**Fig. 2F**), which suggests that direct Mediator recruitment is partially sufficient for generating MpARF1 biological function.

### Partitioning of co-factors into competing ARF clusters

MED15 in many species contains large intrinsically disordered regions (IDRs) with large PrD regions (*34–36*). We used FINCHES (*37*) to predict IDR-IDR interactions between MpMED15 and MpARF1. This supported a strong PrD-PrD interaction relying on MpARF1^F2^ and MpMED15 PrDs (**Fig. 3A**). Predictions using MpARF2 showed a low propensity of MpMED15 interaction for its IDR (fig. S7). The occurrence of PrD and IDRs is a signpost for potential phase separation in proteins (*38, 39*). Indeed, when imaging a line with two-color knock-in fluorescent fusions for MpARF1 and MpARF2 (*40*) at high resolution, we observed that both ARFs form largely non-overlapping nuclear punctae in nuclei at physiological concentrations (**Fig. 3B**), which we refer to as ARF clusters. Colocalization analyses on double transgenic lines expressing MpARF1-mNeonGreen and MpARF2-mScarlet3 on a wild-type background further support the non-overlapping presence of these clusters (fig. S8). We next asked if the interactions with MpMED15 and MpTPL occur within clusters by using bimolecular fluorescence complementation assays. We found that MpARF1 interacts with MpMED15 in punctae, and MpARF2 interacts with MpTPL in similar punctae (**Fig. 3C, D**). These interactions were specific, as MpARF1 did not interact with TPL and MpARF2 did not interact with MED15. Furthermore, MpARF2-MpTPL interaction could not be predicted by means of IDR interaction propensities (fig. S9A), suggesting it may not be driven by chemically specific IDR-IDR interactions. By crosslink-mass spectrometry of recombinant MpARF2 and MpTPL proteins, we confirmed that their interaction is mediated by two different TPL-interacting motifs (EAR-like and LFG) present in MpARF2-MR, which engage with two different MpTPL folded interfaces (fig S9B).

**Fig. 3.**
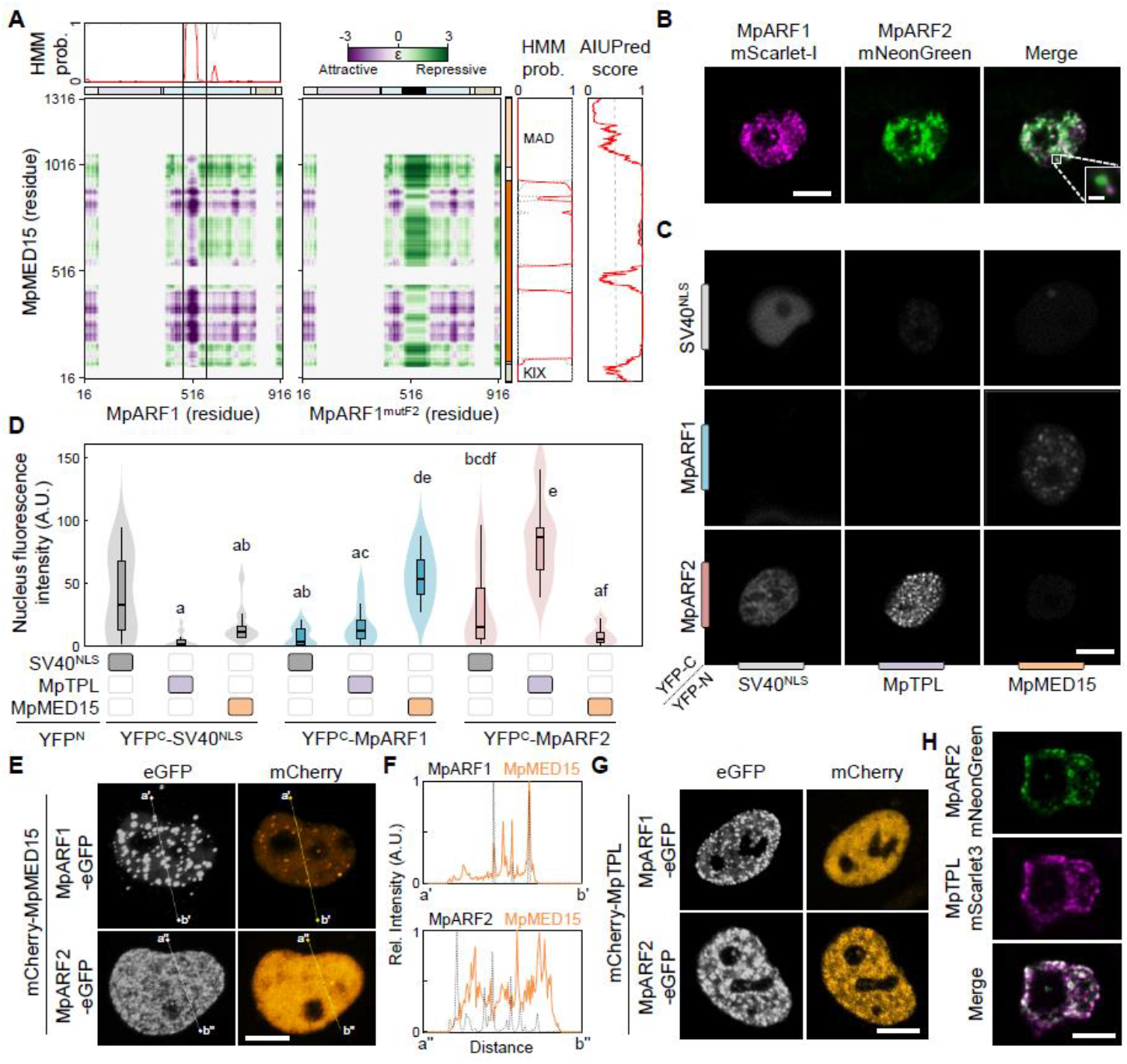
Cofactor partitioning into competing MpARF1 and MpARF2 clusters. **(A)** Prediction of MpARF1-MpMED15 interaction using FINCHES. Attractive and repulsive interactions are plotted onto a protein residue matrix, identifying strong attraction in F2 region of MpARF1 and multiple MpMED15 regions. **(B)** Nuclear punctae of MpARF1-mScarlet-I and MpARF2-mNeonGreen in a double knock-in line (inset shows non-overlap). **(C,D)** Imaging **(C)** and quantification **(D)** of bimolecular fluorescence complementation of a matrix of MpARF1, MpARF2 and SV40NLS control with MpTPL, MpMED15 and SV40NLS control. **(E)** Colocalization of mCherry-MpMED15 with MpARF1-eGFP (top), but not MpARF2-eGFP (bottom) in HeLa cell nuclei. **(F)** Fluorescence intensity along lines drawn in **(E). (G)** Colocalization of mCherry-MpTPL with MpARF2-eGFP (bottom), but not MpARF1-eGFP (top) in HeLa cell nuclei. **(H)** Co-localization of MpARF2-mNeonGreen with MpTPL-mScarlet3 in a Marchantia double transgenic line. Scale bars are 10 µm **(B,C,E,G)** and 0.5 µm the inset **(A)**, and 5 µm **(H)**. In **(D)** boxplots in rainclouds indicate the following parameters: centrum, median; upper bound, first quartile; lower bound, third quartile; whiskers maximum and minimum refer to highest and lowest values, respectively, within 1.5*inter-quartile range (IQR). Statistical groups are determined by Tukey’s Post-Hoc test (p < 0.05) following one-way ANOVA.

This specific partitioning of MED15 and TPL into MpARF1 and MpARF2 nuclear clusters was conserved when protein pairs were expressed in HeLa cells (**Fig. 3E-G**, fig. S10), suggesting that no other plant-specific factors are required. While native co-localization analysis of MpARF1 and MpMED15 was hindered by technical limitations in generating fluorescent MpMED15 fusions, we found that MpARF2 and MpTPL strongly co-localized *in vivo* (**Fig. 3H**), clearly distinct from the poorly overlapping MpARF1 and MpARF2 patterns (**Fig. 2B**, fig S8), and supporting the view that MpARF1 and MpARF2 each recruit co-activators versus co-repressors into competing clusters through direct interactions.

### An evolutionary innovation that switched A-ARFs from repressor to activator

A-class ARFs share the capacity to activate transcription, but it is unclear if this capacity has a single evolutionary origin. Having identified F2 and its mechanism of action, we asked if gain of F2 or part of it may represent the evolutionary innovation that led to auxin-dependent gene activation. Extensive phylogenetic analyses among land plant and algal ARFs demonstrated that PLAAC-detected PrD-like regions are found in the majority of A-class ARFs of all land plants (**Fig. 4A**, fig. S11), except for the seed plant-specific ARF5 subclade (*20*) that plausibly lost the ancestral F2-like region. Additional F2 features, including poly-S/P regions, are also found within the euphyllophyte-specific ARF6/8 subclade, suggesting that F2 is a deeply conserved and ancestral feature (**Fig. 4B**), linked to transcriptional activation in A-class ARFs. If gain of the F2 sequence at the origin of A-class ARFs was causal to the gain of transcriptional activation activity, one would predict that engineering this evolutionary innovation would convert a repressor into an activator. We thus recreated this evolutionary innovation by introducing MpARF1^F2^ into the B-class repressor MpARF2. MpARF1 and MpARF2 middle region sequences are extremely divergent, yet short homology regions between them allowed us to insert either the PrD-region or the whole F2 in homologous locations (fig. S12A). Strikingly, these insertions indeed converted MpARF2 into a transcriptional activator in yeast and plant cells (**Fig. 4C,D**). As predicted, this chimeric MpARF2^A1F2^ protein now directly interacted with MpMED15 in yeast (**Fig. 4E**). In line, colocalization analyses of double transgenic Marchantia lines expressing either MpARF2-mNeonGreen or MpARF2^A1F2^-mNeonGreen and another Mediator subunit, MED16, tagged with mScarlet3 shows a significant increase in Mediator partitioning into MpARF2^A1F2^ clusters (fig S12B). These results are remarkable, as the protein still contains the LFG motif that allows recruitment of MpTPL (*16*), suggesting that the interaction with MpMED15 is dominant over MpTPL interaction. We indeed found this to be the case: interaction of MpARF2^A1F2^ with MpTPL is lost (**Fig. 4E**). Consistent with MpARF2 requirement of MpTPL interaction, MpARF2^A1F2^ was unable to complement the Mp*arf2* loss-of-function mutant, similar to the loss of the entire MR (**Fig. 4G**). Conversely, while loss of the MR or F2 severely hampered the ability of inducible (Glucocorticoid Receptor-fused; GR) MpARF1 protein versions to restore the growth and auxin response of the Mp*arf1* mutant, and replacing the MpARF1-MR for that of MpARF2 (MpARF121) led to a completely inactive protein (fig. S13), as shown before (*16*), integrating ARF1-F2 in the ARF2-MR context (MpARF12^A1F2^1) did partly recover MpARF1 functionality.

**Fig. 4.**
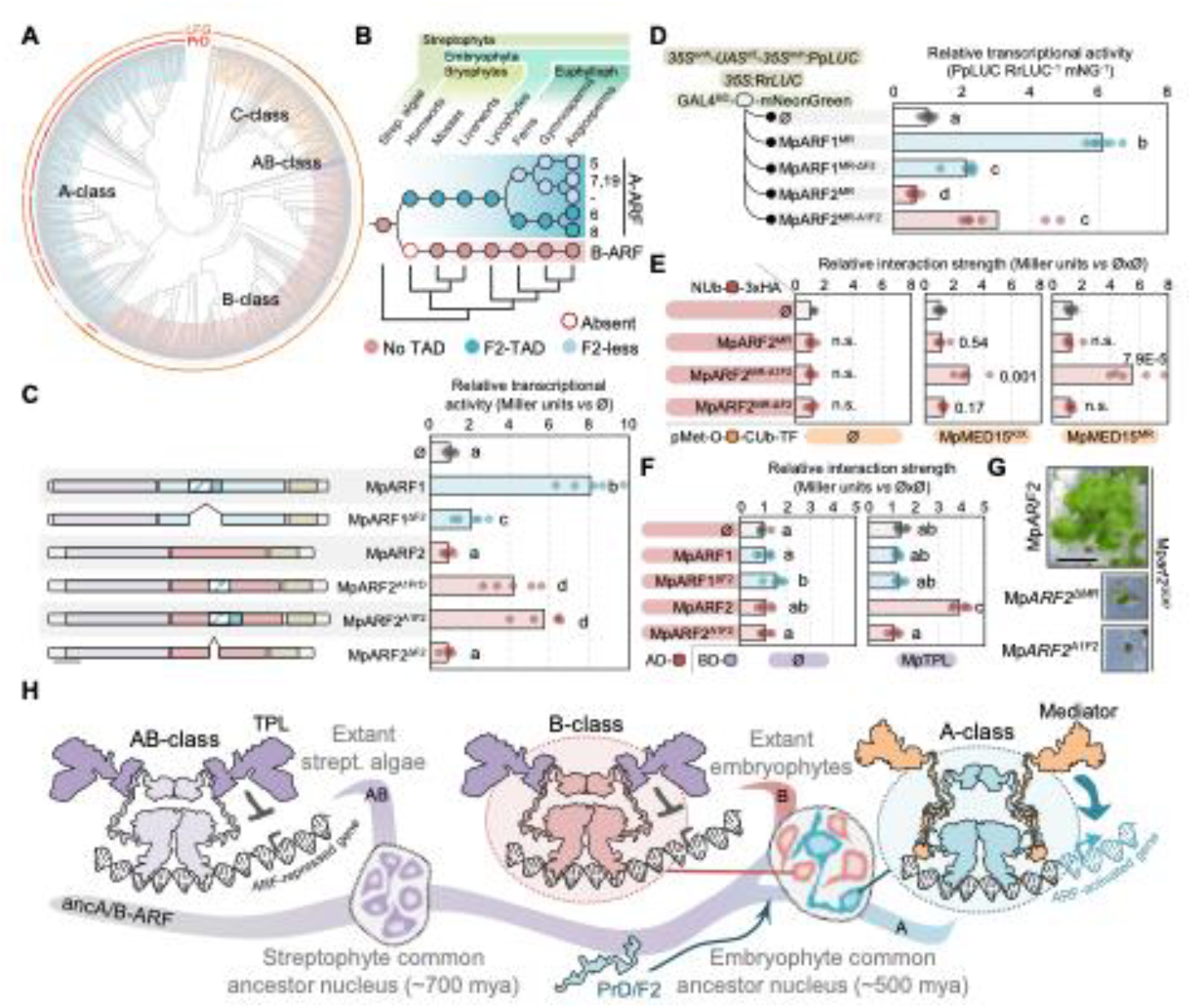
The evolutionary origin of transcriptional activation by A-ARFs. **(A)** ARF phylogeny (classes are indicated) with the presence of the PrD in the middle region, as well as the LFG motif indicated on the periphery. Note that LFG is universally present, and PrD is present in all A-ARFs except a small cluster. **(B)** Reconstruction of the evolutionary trajectory of the F2-TAD showing early origin in the founder of A-ARFs, and loss in a vascular plant subclade. **(C)** Yeast one-hybrid activation assay as a measure of β-galactosidase activity (Miller units) shows that introducing the MpARF1 F2 into the homologous position of MpARF2 converts it into an activator. **(D)** MpARF2 with MpARF1-F2 activates transcription in an Arabidopsis protoplast transcriptional activation assay. **(E)** Yeast cytosolic split-ubiquitin interaction assays on a series of MpARF2 chimaeras shows that MpARF1-F2 introduction into MpARF2 leads to direct interaction with MpMED15. **(F)** Yeast two-hybrid assay on MpARF2 chimaeras shows that MpARF1-F2 introduction causes loss of interaction of MpARF2 with MpTPL. **(G)** Complementation of the Mparf2GE#7 loss-of-function mutant with wild-type MpARF2, MpARF2 lacking its MR and MpARF2 with MpARF1-F2 shows that gain of F2 causes loss of MpARF2 function. **(H)** Schematic model of the A- and B-class divergence driven by the acquisition of an activation domain-capable fragment leading to differential partitioning within the nucleus in land plants. Scale bar, 1 cm (G). In **(C,D,F)**, statistical groups are determined by Games-Howell Post-Hoc test following Welch’s ANOVA (p < 0.05). In **(E)**, statistical differences are determined after log-transformation of data followed by pairwise Welch’s t-test with Holm correction for multiple testing (p < 0.01).

We conclude that F2 is a functional transcriptional activation domain in A-class ARFs, and its acquisition by the ancestral A-class ARF likely marks the evolutionary event that converted the ancestral A/B-ARF transcriptional repressor into a transcriptional activator, overriding the prior repressive function.

## Discussion

Auxin-dependent gene regulation hinges on the antagonistic activity of A-class and B-class ARF transcription factors (*16*). Mutants affecting either class cause dramatic developmental and growth phenotypes (*25, 41*). The biochemical mechanism underlying ARF-mediated gene repression depends on recruitment of the TPL co-repressor (*15, 42, 43*). It has so far remained unclear how A-class ARF’s activate transcription, aside from contrasting evidence among divergent A-class subclades (*21–23*). Furthermore, as our recent reconstruction of the evolutionary history leading to the current auxin response system revealed, A- and B-class ARFs derive from a common repressive A/B-class precursor (*20*). However, activation is essential for auxin-mediated transcriptional response mechanism: auxin destabilizes the Aux/IAA co-repressors that otherwise bind A-ARFs and counteract their function (*15*). This requires Aux/IAAs to act as repressor-activator switches, and, therefore, the evolution of activator ARFs enabled the system’s logic. Hence, a key question is how transcription activation evolved.

Here we discover the mechanism of MpARF1-dependent activation and its evolutionary origin. We first delineate the minimal region required for gene activation by the Marchantia A-class ARF, MpARF1. We discover F2, a PrD-containing region that endows transcriptional activation activity by recruiting the Mediator complex through its MED15 subunit via PrD-PrD interactions. While there are only a few studies on MED15 in plants (*35, 44*), it is the most common co-factor in Mediator-dependent co-activation in other eukaryotes, such as yeasts and mammals (*45, 46*). We show that the F2 region is essential for the function of MpARF1, especially for transcriptional activation, however, our data also suggests additional activation-independent functions within the MR. Consistent with this, previous studies indicate that the Mp*arf1* mutant is devoid of auxin transcriptional responses (*47*), including downregulation of genes such as Mp*YUC2*, which could be related to these additional functions. Interestingly, A-class ARFs retained their LFG TPL recruitment motif within the MR (*19*), and this motif can mediate repression when isolated (*16*). It is possible that this interaction allows for context-dependent repression by A-class ARFs, in an F2 and activation-independent manner.

In addition, we find that Marchantia ARF1 and ARF2 both form discrete clusters within nuclei in lines that carry genomic fluorescent tag knock-ins. These plants are haploid and show normal phenotypes unlike plants that have increased or decreased ARF1 or ARF2 levels (*40*). Thus, these punctae, which we have also found to form *in vitro* as nanoclusters with purified MpARF2 protein (*48*), form at endogenous protein concentrations. While ARF assemblies have been described before in the cytosol (*49, 50*), we interpret these structures to be nuclear clusters, and provide evidence that these are enriched in Mediator complexes (MpARF1) or MpTPL (MpARF2). These new structures may represent transcriptional clusters (in some cases referred to as condensates) involved in RNA polymerase recruitment and activation, or in repression maintenance as found in other eukaryotes (*51, 52*). The most plausible scenario is that the ancestral A/B-ARF had the capacity to form nuclear clusters with ancestral TPL proteins, while the integration of the F2 –or a PrD alone within a serine/proline rich region– in the ancestral land plant A-ARF allowed MED15 (and the Mediator complex) to be partitioned into these clusters and start promoting transcriptional activation. At present, we cannot test the significance of cluster formation, but it is likely that the high local ARF concentrations allow efficient interaction with TPL or MED15 through avidity-based low-affinity interactions (*39*), thus competing with other TF clients for these generic co-factors. In addition, these clusters endow the ARF protein with unique DNA-binding properties (*48*).

Importantly, the insertion of this MED15-recruitment motif converts a repressor (MpARF2) into an activator and thus acts in a dominant manner. We find that the F2 region represents a conserved and ancestral condition in all land plant A-class ARFs, except the euphyllophyte-specific clade (ARF5/7/19). Thus, the insertion of the F2 region in the ancestral A-class ARF gene from an unknown donor sequence, or through other mutational processes such as CAG-strand slippage leading to polyglutamine tract accumulation (*53, 54*), likely converted the A-class into activators, suppressing its repressor capacity.

Our work offers a biochemical mechanism for the transcriptional regulation underlying auxin response, and identifies a differential, competitive chromatin-based co-factor partitioning principle at the heart of the system. It also reveals an evolutionary scenario for the emergence of two antagonistic transcription factors from a common ancestor, pointing to a broader principle in the emergence of transcriptional activation and evolution of transcriptional control.

## Acknowledgements

pMetOYC-DEST (Addgene #105082) and pNX32-DEST (Addgene #105083) were gifts from Christopher Grefen. THY.AP4 and THY.AP5 yeast strains were obtained from the Arabidopsis Biological Resource Center (ABRC). We are also grateful to Roberto Solano for kindly sharing the pEN207 MpMED25 plasmid. This paper was typeset with the bioRxiv word template by @Chrelli: www.github.com/chrelli/bioRxiv-word-template

## Author contributions

Conceptualization: JHG, DW

Data Curation: JHG, MR, DP

Investigation: JHG, MDA, MdR, MGB, MR, ZB, HS, JWB

Formal analysis: JHG, MDA, MdR, MGB, MR, ZB, HS, DP, ASH, HOL

Funding Acquisition: JHG, JWB, ASH, HOL, DW Supervision: ASH, HOL, DW

Validation: JHG, MDA, MdR, MR, ZB, HS, DP, ASH, HOL

Visualization: JHG, MDA, MdR, ZB, HS, ASH Project Administration: JHG, DW

Writing – Original Draft: JHG, DW Writing – Review & Editing: All authors

## Materials and Methods

### Transcriptional activation domain prediction

Initial prediction for activation domains in the MpARF1 primary sequence were done using the TADA predictor (*24*), a neural network tool developed to identify plant activation domains. The analysis was run on the Colab notebook https://colab.research.google.com/drive/1g4tkklihI-dIZV5BEHpPgpqtvofw1BFV. Residues with TAD score above 0.4 are treated as plausible TAD-bearing.

### Molecular cloning and plasmid construction

Oligonucleotides and plasmids used and generated in this study can be found in **Data S2**. Plasmids bearing Marchantia full-length ARF coding sequences derived from cDNA were previously generated (*20*) and used directly or as template for additional cloning purposes. Gateway^TM^ entry vectors were obtained by transferring PCR-amplified sequences into a linearized pEN207 (amplified with JHG079/080) via NEBuilder® HiFi DNA assemblage (New England Biolabs) unless specified. Entry vectors were transferred into the corresponding destination plasmids via LR Clonase II (Invitrogen) reaction for yeast assays and Marchantia complementation assays. Gateway destination plasmids were readily available or build using existing plasmids as templates; pHKDW046-GR was built using pHKDW046 (*16*) as template to introduce a C-terminal Glucocorticoid Receptor fragment, while pHKDW031 and pHKDW037 were built using pMpGWB307 (Addgene #68635) and pMpGWB301 (#68629), respectively, as template to introduce a 3882 basepair-long Mp*ARF2* promoter including the 5’UTR region. ARF fused to sYFP2-TurboID plasmids were obtained by combining the *sYFP2* gene sequence derived from pPLV18 (*55*) and a newly designed plant (Arabidopsis) codon-optimized *TurboID* encoding sequence, based on (*56*), into a pEN207-MCS-sYFP2-TurboID plasmid and inserting MpARF CDS in the cloning site (MCS) via NEBuilder® HiFi DNA assembly reactions. Non-Gateway plant destination plasmids were produced using HiFi reaction with designed oligo fragments. For *Mparf1;Mpncarf* double mutant transformations, a new geneticin (G418) backbone, named pMpGWB500, was built using pMpGWB100 (Addgene #68062) as template replacing the *HPTII* cassette with the *p*Mp*EF1α*-driven *NPTII* cassette from the OpenPlant kit (*57*). pMpGWB100 and pMpGWB500 plasmids bearing mNeonGreen-V5 and mScarlet3-3xFLAG were built as empty vectors for downstream applications and are available upon request. Marchantia *ncARF* editing plasmids were previously described (*58*). For human cell (HeLa) transfection, wild-type sequences of MpARF1, MpARF2, MpMED15, and MpTPLwere inserted into a pcDNA5 mammalian expression vector via Gibson Assembly® (New England Biolabs) adding either a C-terminal eGFP (MpARF1-2) or N-terminal mCherry (MpMED15, MpTPL) sequences in frame.

### Plant Material and Growth Conditions

The Marchantia *ARF* mutants Mp*arf1-4*, Mp*ncarf* and Mp*arf2*^*GE7*^ in the Takaragaike-1 accession (Tak-1; male) had been previously described (*19, 25, 41*). *M. polymorpha* plants were cultured on half-strength Gamborg’s B5 pH 5.5-5.8 with 1% agar at 22°C and constant white light (60 μmol m^−2^ s^−1^), and maintained through asexual reproduction in axenic conditions. A list of all plant lines used in this study can be found in **Data S2**.

### Marchantia transformation and phenotyping

*Marchantia polymorpha* transformants were obtained by an adapted agrobacterium-mediated thalli transformation protocol (*20*) based on (*59*). Briefly, one-to four-week-old plants were transferred to 0M51C medium, fine-chopped with a sterile razor to small pieces, and transferred to 6-well plates. *Agrobacterium tumefaciens* strain GV3101 carrying the appropriate plasmids were inoculated into each well for co-culturing for three days. Washing and plating was performed as the original protocol. Regenerated lines were confirmed by PCR in G1 plants, and, in the case of CRISPR-generated mutations, by sanger sequencing in G1 and G2 plans. To obtain comparable lines expressing different protein versions, Marchantia lines with fluorescently-tagged ARFs were imaged using a Leica SP8X-SMD confocal microscope equipped with a hybrid detector and a pulsed white-light laser (WLL). Citrine was imaged using a water immersion 20X objective (Exc 514 nm, Em 521 – 569 nm). Z-stacks were obtained for each gemma and two or more lines were chosen based on nuclear fluorescence. Phenotyping was performed growing gemmae in medium supplemented with mock (DMSO), and 3 µM 1-naphthyl acetic acid (NAA) or indole-3-acetic acid (IAA) as previously (*16*). Glucocorticoid-inducible ARF assays were performed supplementing the medium with mock or 1 µM dexamethasone. Thallus area was quantified in thallus halves from independent-growing apical notches after 10 days as described before (*60*).

### Yeast transactivation assay

Gal4-DNA Binding Domain (BD) fusions were made by transferring the indicated ARF fragments into the pGBKT7-GW (*61*) vector from pEN207 entry vectors (see above) via LR Clonase II (Invitrogen). Final GBKT7 plasmids were transformed into the Y2HGold strain and selected on SD medium lacking tryptophan (Trp) using the Frozen-EZ Yeast Transformation II Kit (Zymo Research). Quantitative transactivation tests were performed as previously described (*20*). Briefly, GBKT7-bearing Y2HGold haploid strains were mated with a Y187 haploid strain bearing a disarmed plasmid conferring leucine selection, and selected in SD medium lacking leucine, Trp and uracil. Resulting strains were grown in liquid SD lacking Leu, Trp, and Ura and β-galactosidase activity quantification performed as described (*35*) using the substrate ortho-nitrophenyl-β-galactoside (ONPG) for colorimetric measurements.

### Dual luciferase transactivation assays in Arabidopsis protoplasts

Final effector constructs were made by transferring ARF sequences from entry vectors to the pMON-Gal4DBD-GW-mNeonGreen plasmid (*20*) via LR Clonase II (Invitrogen). A previously available pGreenII 5xGal4 UAS dual luciferase construct was used as reporter (*35*). Assays were performed by co-expressing the effector plasmids and the reporter plasmid in Arabidopsis (Col-0) leaf mesophyll protoplasts as previously described (*62*). Briefly, *Arabidopsis* young leaves were harvested and the epidermal layer was removed using magic tape. Exposed mesophyll cells were released by incubating the leaves with 1% cellulase and 0.2% macerozyme. Protoplasts were isolated and transfected with equal amounts of effector and reporter plasmid via polyethylene glycol-mediated transfection and incubated for 16 hours. After incubation, protoplasts were collected and lysed using Passive Lysis Buffer (Promega). First, supernatant mNeonGreen-derived fluorescence was measured using a BioTek Synergy H1 Multimode Reader (Agilent) as a protein-level approximated quantification. Next, luciferase activities in the same supernatants were quantified with the Dual-Glo Luciferase Assay System (Promega) using the same reader. Ratio of Firefly luciferase/Renilla luciferase luminescence was normalized against the fluorescence measurements to obtain transactivation activity. Each supernatant was measured three times, and at least four independent transfection events were used per effector to obtain statistical differences between effector activities, as specified in the figure captions.

### Proximity labelling

Biotin proximity labelling was performed as previously described with modifications (*63*). Briefly: ten-day-old *Marchantia* gemmalings were treated with 50 µM biotin (B4501; Sigma) for 24 hours. The following day gemma were harvested and grinded in liquid nitrogen to a fine powder. Powder was suspended in nuclei isolation buffer (20 mM Tris (pH 7.5), 25% glycerol, 20 mM KCl, 2 mM EDTA, 2.5mM MgCl_2_ 250 mM Sucrose,1 mM DTT and 1 mM PMSF) and agitated on a rotor for 20 minutes at 4°C. Homogenate was filtered through a 100 µm nylon mesh and spun down for 15 min at 3000 RPM at 4°C. Crude nuclei were lysed in RIPA buffer (50mM Tris pH 7.5-8.0, 150 mM NaCl, 2 mM MgCl_2_,1 mM EDTA, 1 mM DTT,1% NP40, 0.5% Sodium Deoxycholate, 0.1% SDS, 10% Glycerol, 1x Cocktail protease inhibitor). Lysis was aided by sonication using a Qsonica Q500 waterbath sonicator with 3 pulses of 15/30 seconds on/off, at 100% amplitude. Lysate was spun down for 30 minutes at 21000x*g* at 4°C. Excess biotin in lysate was removed using 10 kDa Amicon filters. Lysate was added to filters and centrifuged for 30 minutes at 5000x*g* at 20°C. Buffer was exchanged by adding 1 ml of RIPA buffer and centrifuged again for 30 minutes at 5000x*g* at 20°C. Buffer exchange was repeated for a total of three times. Retinate containing the proteins was transferred to new tubes and protein content measured using the BCA assay. Equal amount of protein (∼2 mg) was added to 50 µl pre-equilibrated Streptavidin Mag Sepharose® beads and rotated overnight at 4°C. The following day beads were washed trice with 1 ml RIPA, trice with 1 ml RIPA without detergents and trice with 1 ml 50 mM ammonium bicarbonate (ABC). After the last wash acrylamide in 50 mM ABC was added to a final concentration of 50 mM and incubated for 30 minutes at room temperature. After alkylation, beads were washed once with 50mM ABC and incubated with 0.5 µg trypsin in 50 mM ABC overnight at room temperature. Following day peptides were acidified and desalted using homemade C18 µcolumns.

### Protein production and crosslinking mass spectrometry

The MpTPL CDS was introduced into PCR-amplified pET His6 MBP TEV LIC plasmid (Addgene #29656) via HiFi DNA reaction and the protein produced as previously with minor modifications (*17, 20*). The 6xHis-MBP-tagged TPL fusion protein was expressed in *E. coli* strain Rosetta 2(DE3) (Novagen) by inducing with 0.3 mM IPTG for 16 hours at 20 °C. Cell-free extracts were used to purify proteins through two consecutive affinity chromatographies (1^st^: HisTrap^TM^ High Performance, 2^nd^: MBPTrap^TM^ High Performance; Cytiva) followed by a size exclusion chromatography (Superdex^R^200 Prep Grade, Cytiva) using an ÄKTA Pure 25 system. MpARF2 full-length protein was produced as previously (*48*). To identify interaction surfaces through crosslink, recombinant MpARF2 and MpTPL were mixed in a 1:1 ratio and reacted with bis(sulfosuccinimidyl)suberate (BS3) in a 1:0.5 ratio (protein:crosslinker) as previously described (*64*), then subjected to mass spectrometry.

### Mass spectrometry and data processing

Proximity labeled and crosslinked samples were measured on a Thermo Exploris 480 coupled to a Vanquish Neo nano LC as previously described (*63*). Raw mass spectrometry data is deposited on MassIVE (MSV000101542) and Pride (PXD077484).

Thermo RAW files were searched using MSFragger v21 (*65*) using the built in LFQ-MBR workflow against the *Marchantia polymorpha* proteome (UP000077202). MSFragger combined_protein.tsv output file was analyzed using the DEP/DEP2 functionalities in R (*66*). Data was filtered to have at least three valid values in at least one group, normalized using vsn and imputed using a mixed imputation approach with knn used for valugroup,ing at random and QRLIC for values missing not at random. Statistically significant enriched proteins were determined using limma against Tak-1 as control.

For crosslinked samples RAW Thermo MSMS files were converter to mgf file format using MS Convert (*67*). Next xiSearch (v1.8.1) (*68*) was used to identify crosslinked peptides. The database contained the protein sequences of MpARF2 and MpTPL without enrichment tags and the reverse sequences as decoys. Search parameters were as follows: MS tolerance, 6 ppm; MS/MS tolerance, 10 ppm, enzyme, trypsin/P, missed cleavages, 2, crosslinker, BS3; fixed modification, carbamidomethylation of cysteine; variable modification, oxidation of methionine, deamidation of asparagine and glutamine and modification by BS3 (BS3, BS3Amidated, BS3Hydrolized) with BS3 reaction specificity at lysine, serine, threonine, tyrosine, and N termini of proteins for the NHS-ester group. Residue pair false discovery rate (FDR) was estimated using xiFDR (v2.2.1) (*69*) with an acceptance of 5%.

### Yeast cytosolic split-ubiquitin and yeast-two hybrid assays

Cytosolic split-ubiquitin assays were adapted from (*70*). Briefly, baits carrying a carboxy-terminal synthetic TF and the C-end ubiquitin fragment and an amino-terminal mOST4 membrane-tagging fusions were generated via LR Clonase II (Invitrogen) recombination from entry plasmids into the pMetOYC-Dest plasmid (Addgene #105082). Prey were produced identically by fusing the N-terminal moity of ubiquitin to the amino-end of the gene of interest using the pNX32-Dest (Addgene #105083) as destination plasmid. Yeast strain THY.AP5 was transformed with pNX32-derived expression vectors, strain THY.AP4 was transformed with pMetOYC-based bait plasmids, and haploid transformants selected in SD medium without Leu or Trp, respectively. Diploid strains were produced by mating and selected in SD medium lacking Leu and Trp.

Yeast-two hybrid assays were performed as previously (*20*). Bait Gal4-DNA Binding Domain fusions were generated as described above with the appropriate entry vectors transferred to pGBKT7-GW (*61*). Similarly, prey Gal4-Activation Domain (AD) constructs were generated via LR Clonase II (Invitrogen) from the entry vectors into pGADT7-GW (*61*). Strain Y187 was transformed with pGADT7-derived expression vectors, while strain Y2HGold was transformed with pGBKT7 vectors, and selected in SD medium without Leu or Trp, respectively. Diploid strains were produced by mating and selected in SD medium lacking Leu, Trp and Ura.

Quantitative interaction assays in either experimental set up were performed in liquid medium by quantifying β-galactosidase activity as described (*20*).

### RNA isolation, cDNA synthesis, and qRT-PCR analysis

Ten-day old Marchantia gemmalings were collected, flash-frozen in liquid nitrogen and grinded into powder using a bead shaker. Total RNA was extracted with a RNeasy Plant Mini Kit (Qiagen) according to the manufacturer’s instructions including an on-column DNase I treatment (Qiagen). cDNA was prepared from 1 μg total RNA with an iScript cDNA Synthesis Kit (Bio-Rad) with poly-T oligo following manufacturer instructions. Mp*WIP* (Mp1g09500) and Mp*YUC2* (Mp8g08780) expression were analyzed as previously using double reference genes (*20*).

### Protein sequence analyses

In silico analyses for intrinsically disordered regions (IDRs) and Prion-like domain sequence presence were performed using available prediction tools. Either IUPred3 (*71*) with long disorder and medium smoothing, or AIUPred v2 (*72*) with standard parameters were used on Marchantia ARF and ncARF protein sequences to identify IDRs and scores plotted accordingly. PLAAC (*73*) was then employed to identify probable prion sequences in the same sequences using a core length of 60 and a relative weighting of background probabilities (α) of 100. Identification of Mediator subunits in *Marchantia polymorpha* was done using PharaohFUN v2 (*30*) on batch mode with Arabidopsis sequences as queries, and summary can be found in Table S2. To predict intermolecular interactions within IDRs between ARFs and Mediator tail subunits, the deep learning-based framework FINCHES v0.1.3 (*37*) was used.

For species-wide analyses, previously phylogenetically analyzed sequences (*20*) were further subjected to curation to use full-length proteins by means of DBD and PB1 presence. The intermediate sequence was set as the middle region and subjected to PLAAC analyses. COREscore > 1 and/or LRR >30 were used to define PrD-like presence, and binary presence/absence plotted using iTOL v7.5.1. For LFG peptide presence, a class-by-class alignment of these curated sequences was aligned using M-COFFEE from T-COFFEE v13.46 (*74*) and presence determined manually.

### *In vivo* high-resolution confocal microscopy

The nuclear localization of mNeonGreen- and mScarlet-I/3-tagged proteins expressed in *M. polymorpha* was visualized using dormant gemma on a Leica STELLARIS inverted confocal microscope. Images of nuclei from rhizoid initial cells were captured using an 86x objective lens with mNeonGreen (Ex/Em: 488/499-556 nm), and mScarlet-I or mScarlet3 (Ex/Em: 569/584-634) filters. Fluorescence was detected in photon-counting mode with time-gated acquisition to reduce autofluorescence. The acquisition parameters were: 100 hz speed, line accumulation of 3, and system optimized z-stacks to apply deconvolution. To quantify colocalization, the Pearson Correlation Coefficient (PCC) and Manders’ coefficients were calculated using JaCoP v2.0 (*75*) (http://rsb.info.nih.gov/ij/plugins/track/jacop.html) for Fiji (http://fiji.sc) on the deconvolved images. One nucleus per image was selected as region of interest (ROI) and Costes’ automatic threshold was applied to each channel. Significance (*p-*value) was evaluated using the Costes’ approach, included in JaCoP v2.0, generating 200 randomized images.

### Bimolecular fluorescence complementation assays

To perform bimolecular fluorescence complementation assays, entries bearing MpARF1/2 or MED15/TPL coding sequences were recombined into pMDC43-YFC and pMDC43-YFN (*76*), respectively through LR Clonase II (Invitrogen) reactions. Pairs of *Agrobacterium tumefaciens* GV3101 strains containing binary plasmids were used to infiltrate 4-week-old *N. benthamiana* leaves. Two days after infiltration, abaxial epidermal pavement cells of infiltrated leaves were analyzed with a Leica SP5 confocal microscope. Reconstituted YFP signal was detected with emission filters set to 503 to 517 nm. Nuclei presence in cells was verified by transmitted light. Signal intensities were measured by manually defining nuclei area of interest based on transmitted light, masking and quantifying in the yellow fluorescence-captured channel.

### Co-localization analyses in mammalian cells

HeLa cells were cultured in Dulbecco’s Modified Eagle Medium (DMEM) supplemented with 10% fetal bovine serum (FBS) and 1x penicillin-streptomycin at 37°C with 5% CO_2_. Cells were seeded in 35 mm imaging dishes (Cellvis, D35C4-20-1-N) a day prior to transfection. At approximately 70% confluency, cells were transfected with 250 ng of MpARF1- or MpARF2-eGFP and 1000 ng of mCherry-TPL or -MED15 using Lipofectamine 3000 transfection reagent (Life Technologies, L3000008), according to the manufacturer’s instructions. Fresh medium containing 2 µg/ml Hoechst dye was added twenty-four hours post-transfection.

Cells were imaged in an environmental chamber (Oko labs, H301-K-FRAME) maintained at 37°C with 5% CO_2_. Images were acquired using a Nikon Ti-2E microscope equipped with an Andor Dragonfly 200 spinning-disc confocal unit, a 60×/1.4 oil-immersion Nikon objective (MRD71600), and a Zyla 4.2 sCMOS camera following excitation with 488 and 561 nm lasers to detect eGFP and mCherry, respectively. Images were processed and analyzed using FIJI/ImageJ. To assess the colocalization of MpARFs with MED15, fluorescence intensity profiles of 488 nm (MpARF-eGFP) and 561 nm (mCherry-MED15) on three random lines were obtained from maximum-intensity projections. Intensity values were normalized to the maximum intensity per line and analyzed by powers of ten to improve the signal-to-background ratio. The fraction of MpARF peaks overlapping with MED15 over total MpARF peaks is reported. To assess MpARF co-localization with TPL, background subtrated fluorescence of individual z-stack was analyzed. A nucleus was masked as a ROI and Pearson’s correlation coefficient was calculated for each ROI using the Coloc 2 plugin (FIJI/ImageJ).

## Inventory of Supplementary Materials

**Supplementary Figures 1-13**

**Supplementary Tables 1,2**

**Supplementary Data files S1, S2 (separate files)**

**Fig. S1.**
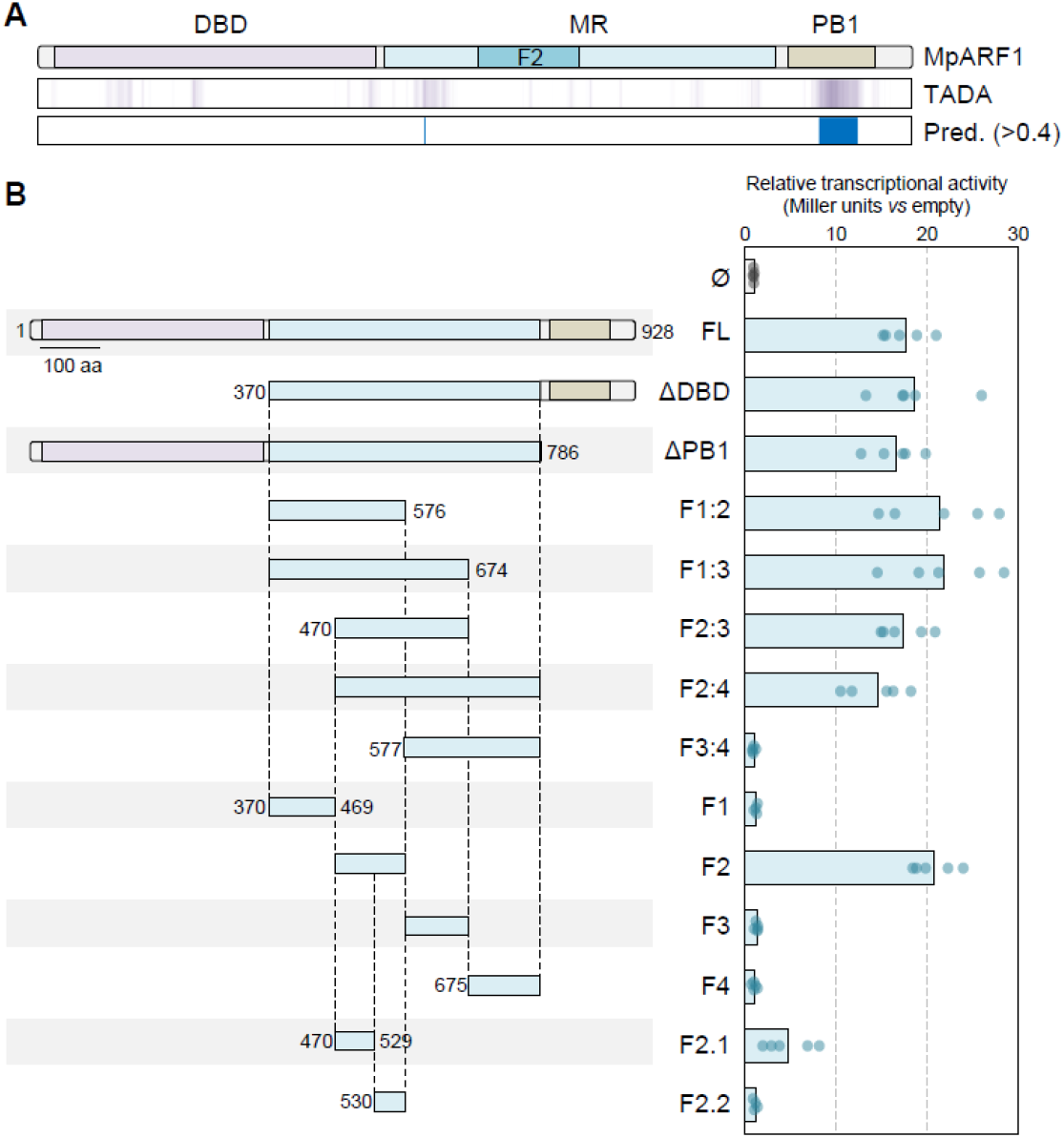
Mapping of MpARF1 transcriptional activation domain. **(A)** TADA prediction of transcriptional activation domains in MpARF1 protein. TADA score is plotted as a minimum (white)-maximum (lilac) color range. Threshold in prediction (Pred., blue) to determine possible TAD has been set in 0.4 (TADA score). **(B)** Yeast-based transcriptional activation assay using MpARF1 and subsequent mapping of a single TAD with this technique.

**Fig. S2.**
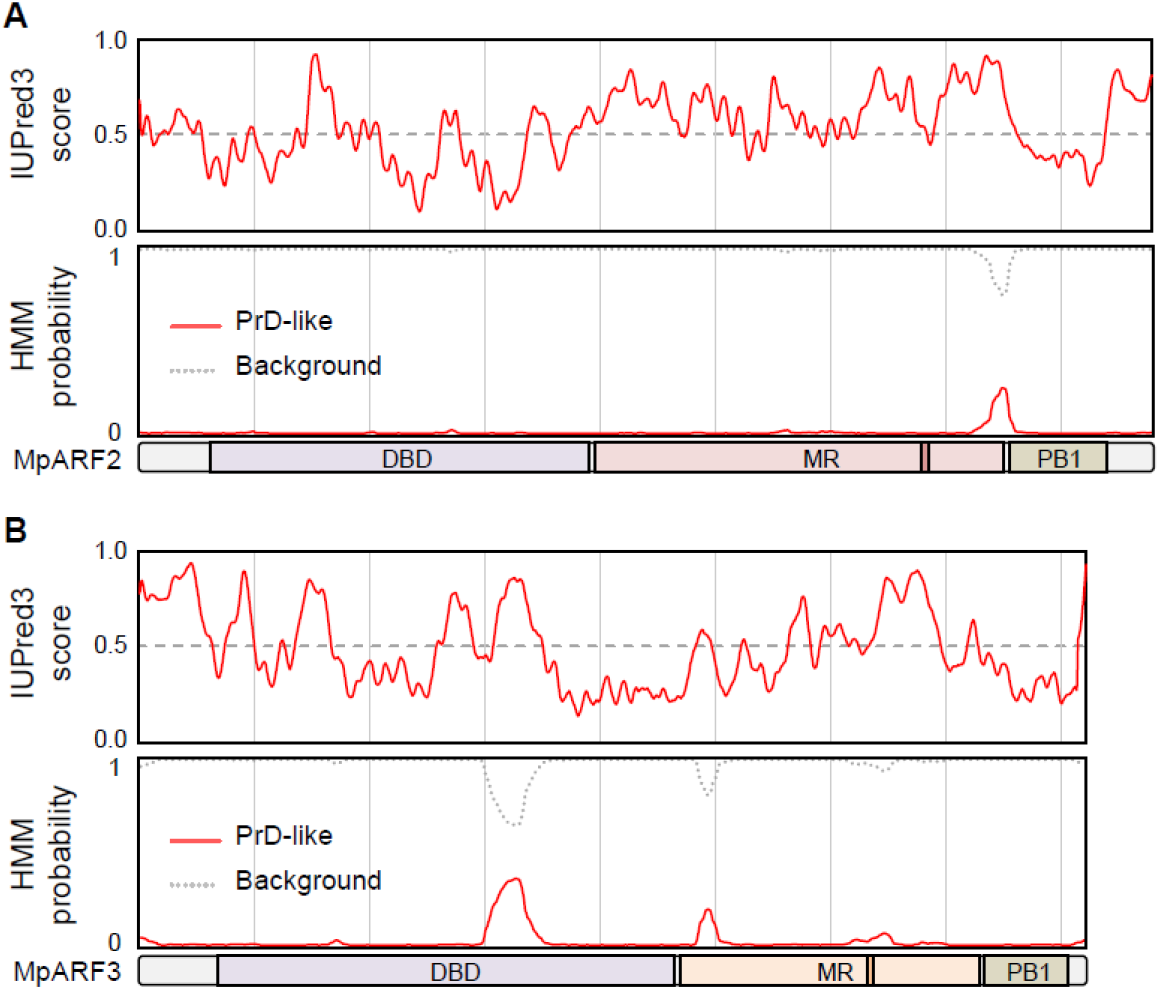
Intrinsic disorder and prion domain-like analyses in Marchantia. IUPred3 and PLAAC results of the B-class ARF, MpARF2 **(A)** and the C-class ARF, MpARF3 **(B)**, showing the intrinsically disordered regions (IUPred3) and the lack of predicted PrD (PLAAC).

**Fig. S3.**
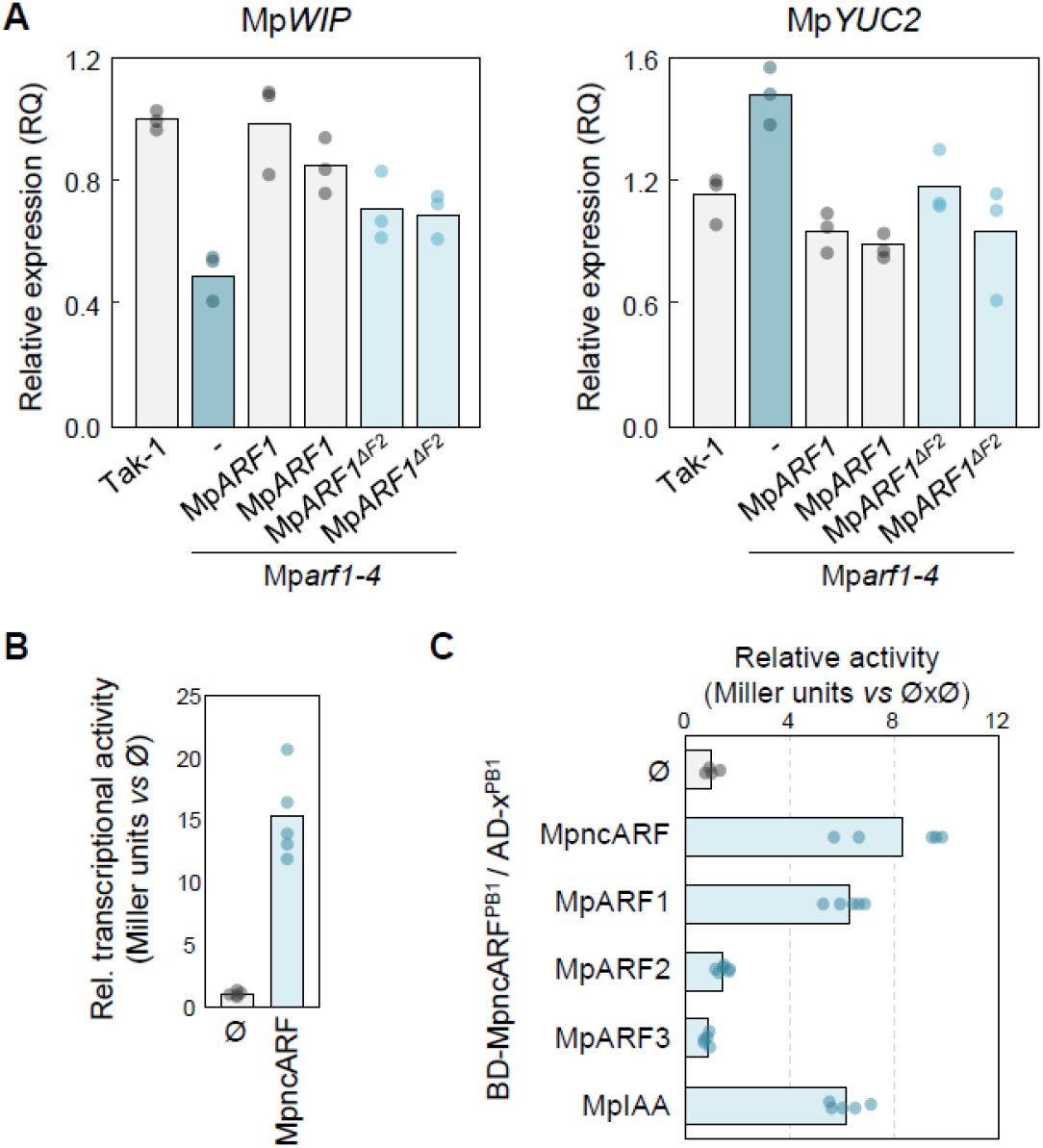
Lack of MpARF1 F2 is likely supplemented by MpncARF. **(A)** Mp*WIP* (Mp1g09500) and Mp*YUC2* (Mp8g08780) gene expression measured by quantitative retrotranscription-PCR. **(B)** Yeast one-hybrid assay showing of transcriptional activation capacity of MpncARF measured by β-galactosidase activity. **(C)** Yeast two-hybrid interaction assay between MpncARF PB1 and the other auxin elements’ PB1s measured by β-galactosidase activity.

**Fig. S4.**
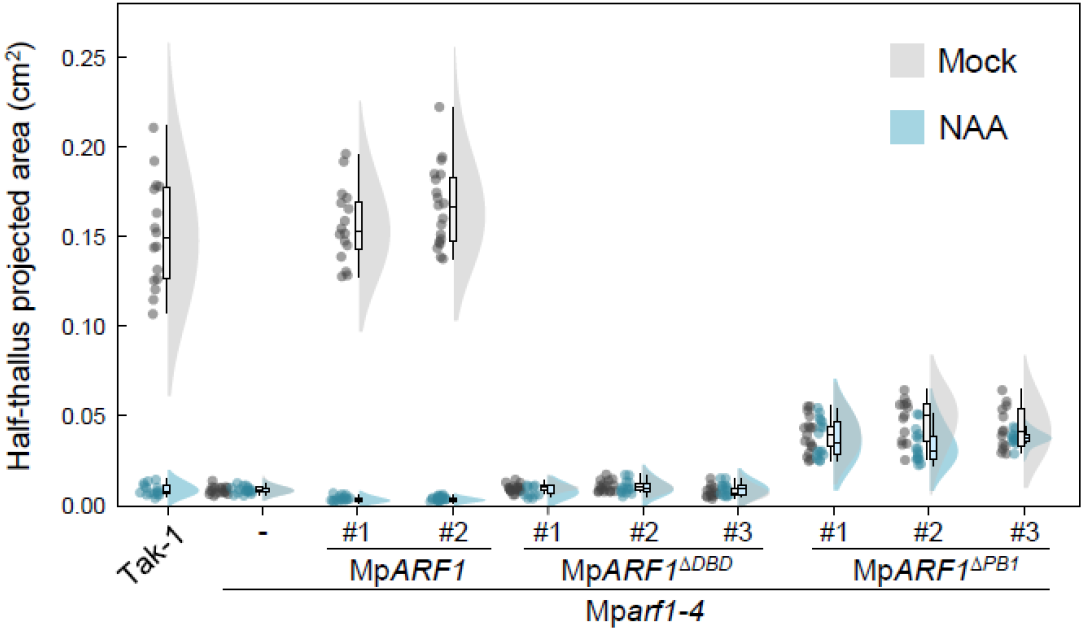
MpARF1 functionality relies in the ability to bind DNA. Mp*arf1-4* complementation assays using MpARF1 and DBD and PB1-deleted versions. The lack of DBD is essential, the lack of PB1 complements part of the function but abolishes response to auxin.

**Fig. S5.**
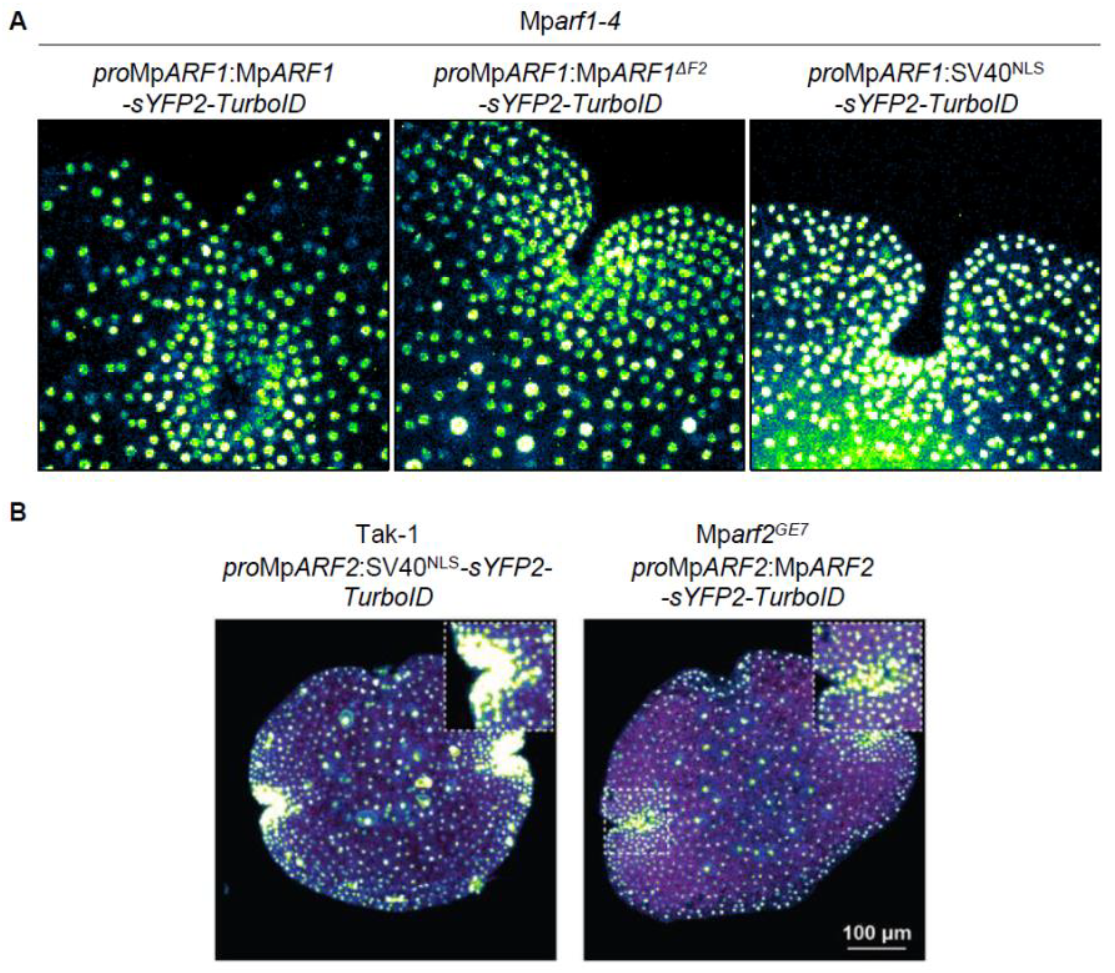
TurboID-tagged MpARFs are functional. **(A)** Localization of sYFP2-TurboID-tagged MpARF1, MpARF1ΔF2 and NLS expressed under a endogenous 4.8 kb Mp*ARF1*promoter in Mp*arf1-4* in the apical notch. **(B)** Localization of sYFP2-TurboID-tagged MpARF2 and NLS expressed under a endogenous 3.9 kb Mp*ARF2* promoter in Mp*arf2GE7* and a wild-type plant (Tak-1), respectively, in dormant gemma and inset of apical notch.

**Fig. S6.**
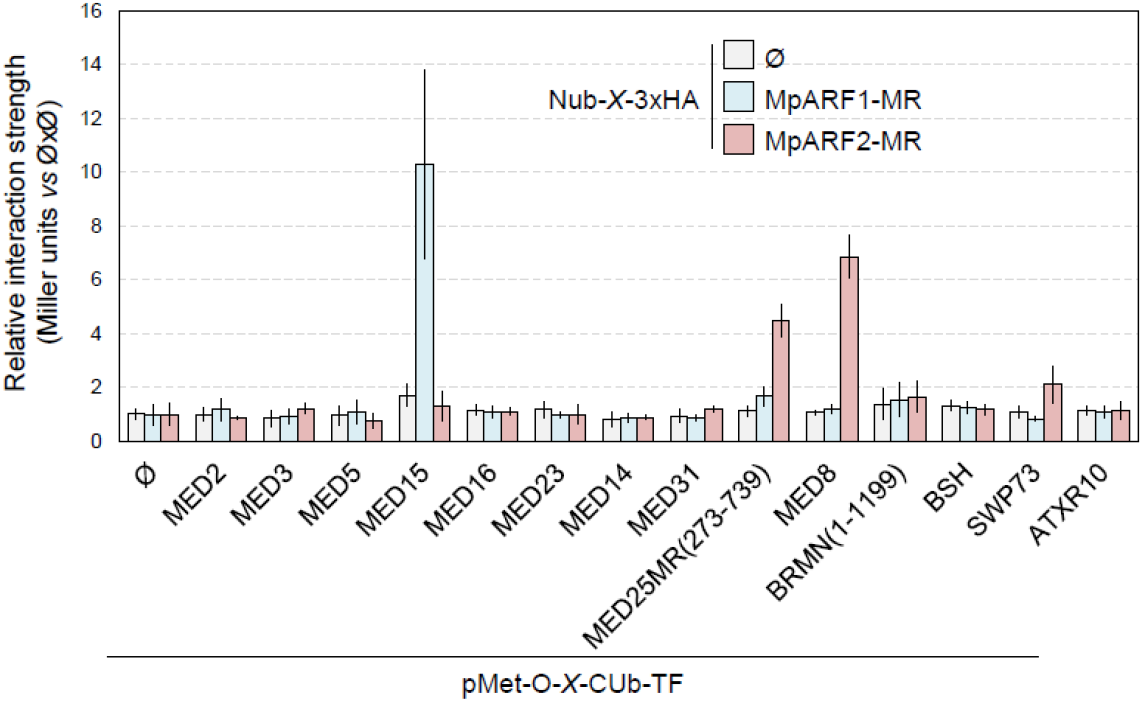
Interaction of MpARF1 and MpARF2 with transcriptional machinery. Yeast cytosolic split-ubiquitin interaction assays between MpARF1 or MpARF2 and Mediator subunits, SWI/SNF2 subunits, and MpATXR10.

**Fig. S7.**
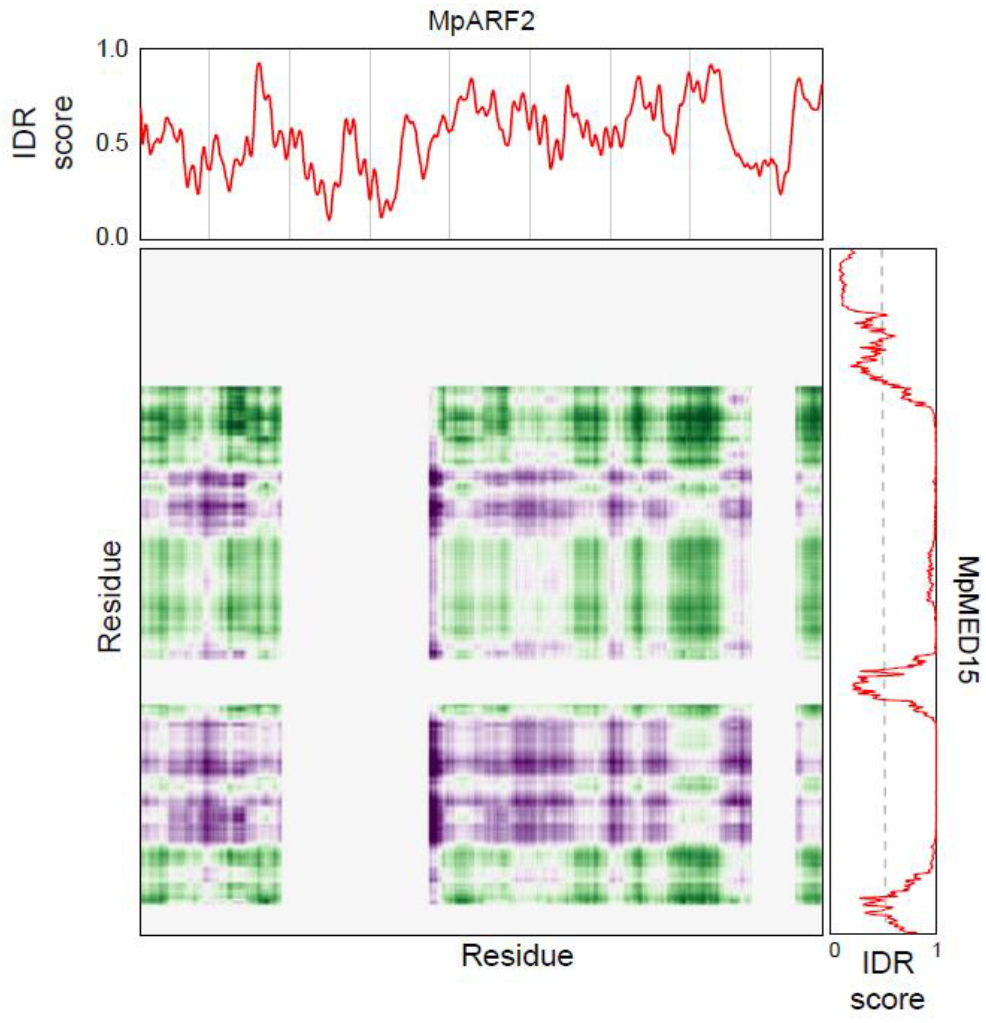
MpARF2 and MpMED15 intrinsically disordered regions do not show interaction hotspots. FINCHES interaction propensity for MpARF2 with MpMED15 full length sequences. Intrinsic disorder prediction is indicated for both proteins. Vertical bars in MpARF2 IDR prediction represent 100 amino acid stretches.

**Fig. S8.**
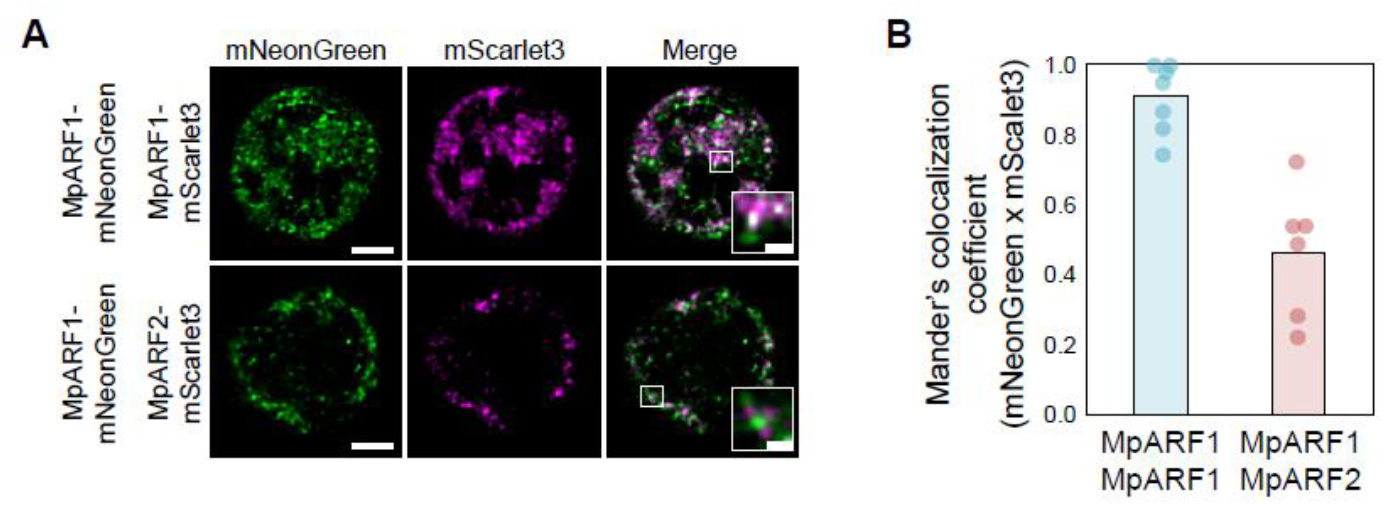
MpARF1 and MpARF2 are differentially distributed in the nucleus. **(A)** High-resolution imaging and colocalization analyses. **(B)** Co-partitioning measurement as Mander’s coefficient between MpARF1-mNeonGreen and a partner, either MpARF1-mScarlet3 or MpARF2-mScarlet3 (see methods). Scale bars are 2.5 µm (0.5 µm for insets).

**Fig. S9.**
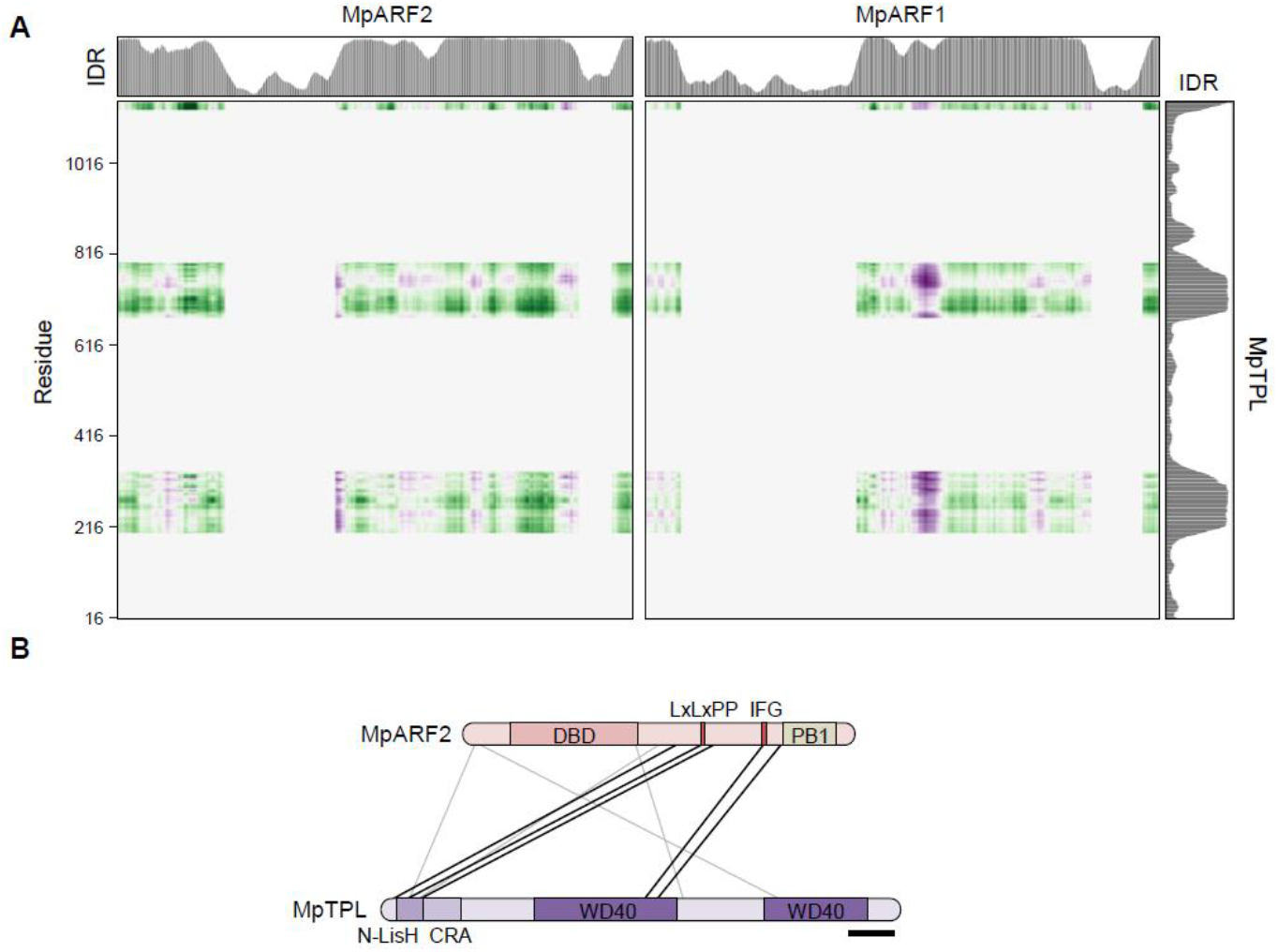
Interaction prediction through intrinsically disordered regions cannot predict MpARF2-MpTPL interaction. **(A)** FINCHES interaction propensity for MpARF2 and MpARF1 with MpTPL full length sequence. Intrinsic disorder prediction is indicated for each protein. **(B)** Schematic representation of direct links between MpARF2 and MpTPL found through crosslink mass spectrometry of recombinant proteins. Black lines indicate significant crosslink between peptides while grey are non-statistically supported. Scale bar = 100 amino acids.

**Fig. S10.**
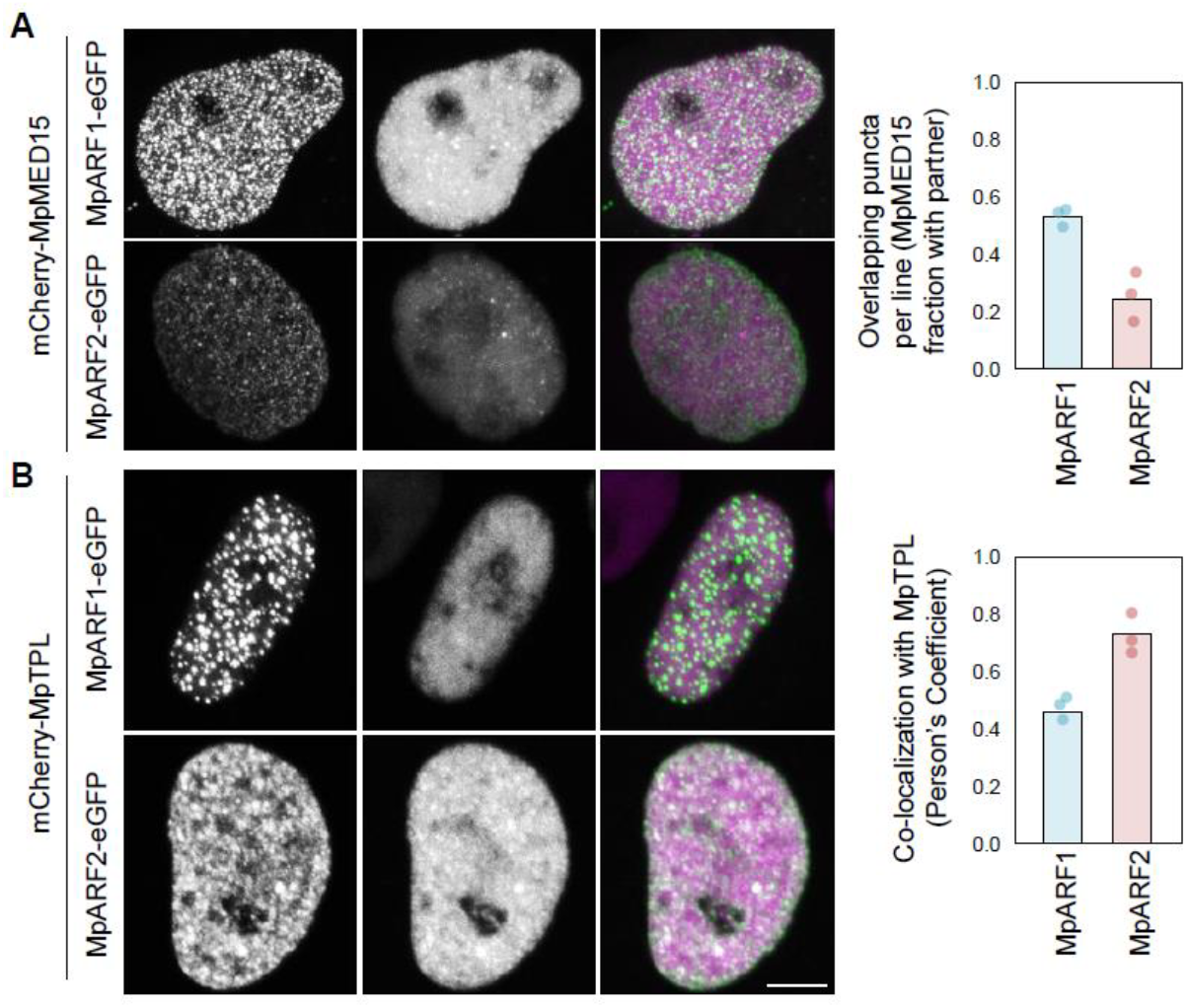
MpARF1 and MpARF2 partition with specific co-effectors. Co-localization of MpMED15 **(A)** or MpTPL **(B)** fused to mCherry with MpARF1 and MpARF2 fused to eGFP in HeLa cells. Quantification of MpMED15 through PCC was not possible due to technical limitations and random line with co-localizing partner fraction was used. Each experiment was run three independent times, using at least ten random nuclei in each. Dots in barplots indicate the average of independent experiments and bar the mean of all pooled. Scale bar = 5 µm; all image at scale.

**Fig. S11.**
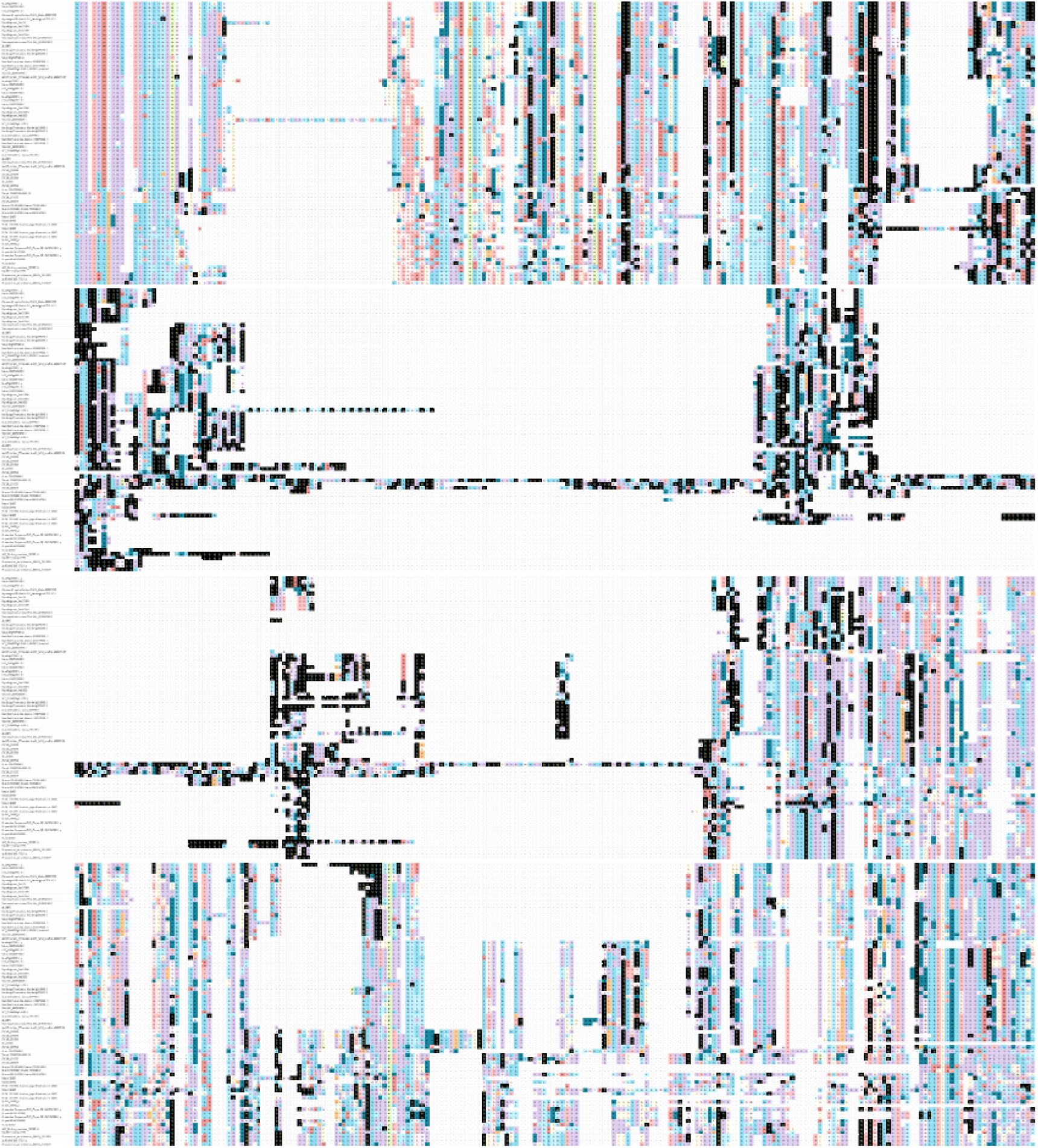
F2 regions are ancestral and conserved in A-class ARFs. Glutamine residues (Q) highlighted in white on with black background. Protein sequence alignment performed with M-COFFEE and manually curated of A-ARF proteins from multiple lineages focused on the middle regions.

**Fig. S12.**
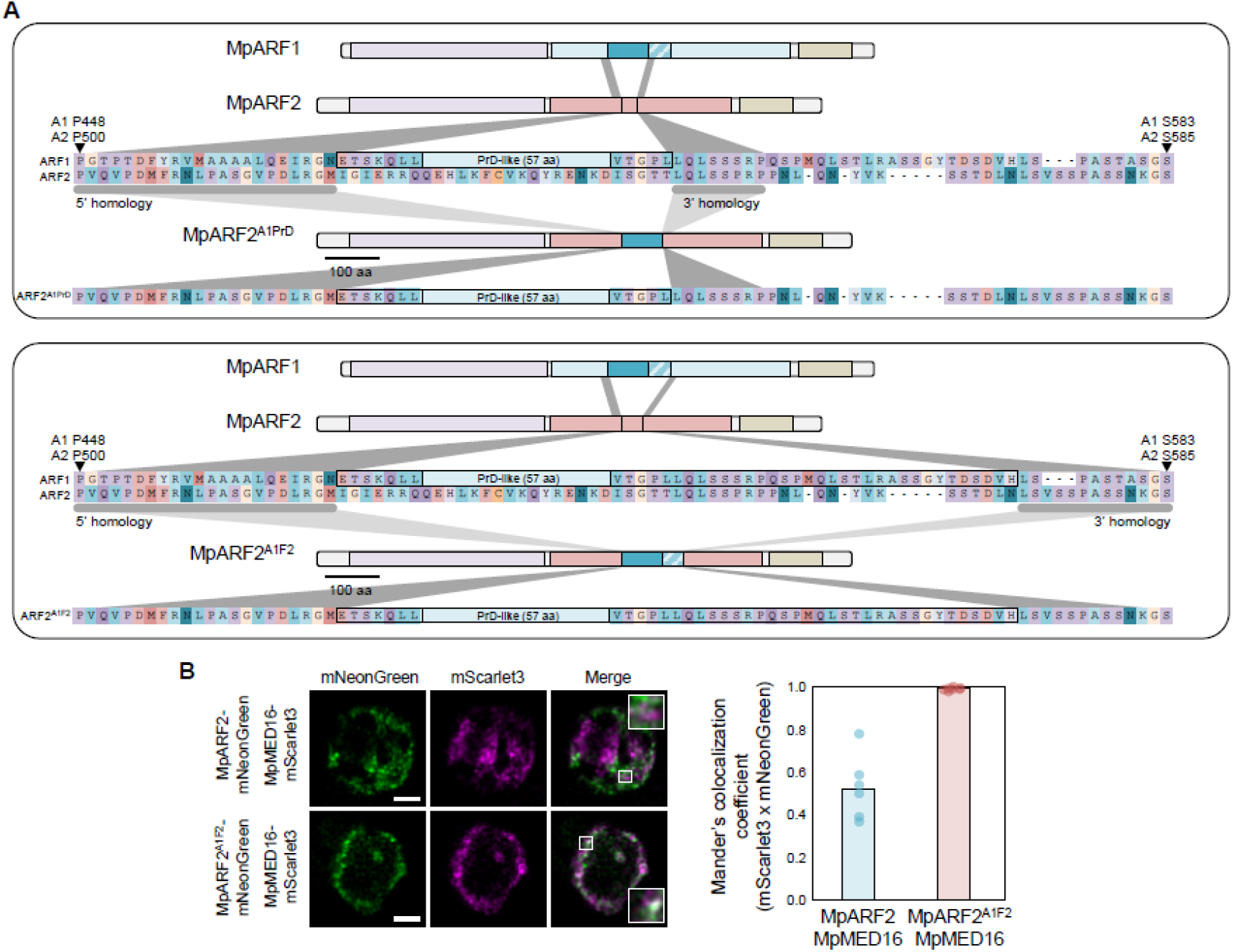
MpARF2 can be converted into an activator. **(A)** Strategy for activation domain introduction into MpARF2 through microhomology regions. **(B)** High-resolution imaging of MpMED16-mScarlet3 coexpressed with MpARF2 and MpARF2A1F2 fused to mNeonGreen (left). Co-partitioning measurement as Mander’s coefficient between MpMED16-mScarlet3 and a partner, either MpARF2-mNeonGreen or MpARF2A1F2-mNeonGreen (right, see methods).

**Fig. S13.**
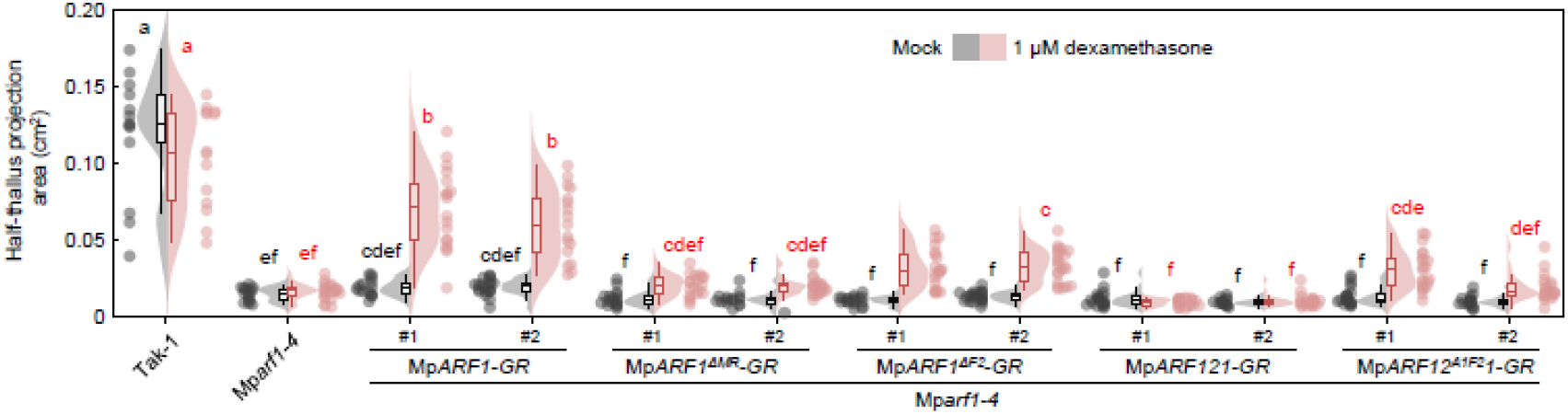
Activator-MpARF2 middle region can rescue MpARF1 native activation function. Quantification of thallus growth in Mp*arf1-4* mutant complemented with dexamethasone (DEX) inducible versions of MpARF1. Introduction of MpARF2 MR in MpARF1 is detrimental for growth, while the addition of F2 allows the version to partly complement the mutant phenotype upon induction. Boxplots in rainclouds indicate the following parameters: centrum, median; upper bound, first quartile; lower bound, third quartile; whiskers maximum and minimum refer to highest and lowest values, respectively, within 1.5*inter-quartile range (IQR). Statistical groups are determined by Tukey’s Post-Hoc test (p < 0.05) following one-way ANOVA.

**Table S1.** Amino acid composition of MpARF domains.

| Amino acid |  |  | Full length |  | MR |  | F2 |  | PrD |  | polyPS |  | DBD |  | PB1 |  |
| --- | --- | --- | --- | --- | --- | --- | --- | --- | --- | --- | --- | --- | --- | --- | --- | --- |
| MpARF1 | Ala | A | 61 | 6.60% | 26 | 6.30% | 1 | 0.90% | 0 | 0.00% | 1 | 2.30% | 28 | 7.60% | 7 | 4.90% |
|  | Arg | R | 63 | 6.80% | 24 | 5.80% | 4 | 3.70% | 2 | 3.50% | 2 | 4.70% | 29 | 7.90% | 9 | 6.30% |
|  | Asn | N | 25 | 2.70% | 13 | 3.10% | 2 | 1.90% | 2 | 3.50% | 0 | 0.00% | 7 | 1.90% | 5 | 3.50% |
|  | Asp | D | 51 | 5.50% | 23 | 5.60% | 3 | 2.80% | 0 | 0.00% | 3 | 7.00% | 18 | 4.90% | 10 | 7.00% |
|  | Cys | C | 11 | 1.20% | 4 | 1.00% | 0 | 0.00% | 0 | 0.00% | 0 | 0.00% | 4 | 1.10% | 3 | 2.10% |
|  | Gln | Q | 86 | 9.30% | 60 | 14.50% | 40 | 37.40% | 36 | 63.20% | 3 | 7.00% | 18 | 4.90% | 8 | 5.60% |
|  | Glu | E | 38 | 4.10% | 10 | 2.40% | 2 | 1.90% | 1 | 1.80% | 0 | 0.00% | 19 | 5.10% | 9 | 6.30% |
|  | Gly | G | 64 | 6.90% | 30 | 7.30% | 2 | 1.90% | 0 | 0.00% | 2 | 4.70% | 21 | 5.70% | 12 | 8.50% |
|  | His | H | 23 | 2.50% | 6 | 1.50% | 2 | 1.90% | 1 | 1.80% | 1 | 2.30% | 15 | 4.10% | 2 | 1.40% |
|  | Ile | I | 25 | 2.70% | 7 | 1.70% | 0 | 0.00% | 0 | 0.00% | 0 | 0.00% | 15 | 4.10% | 3 | 2.10% |
|  | Leu | L | 82 | 8.80% | 39 | 9.40% | 14 | 13.10% | 4 | 7.00% | 8 | 18.60% | 30 | 8.10% | 13 | 9.20% |
|  | Lys | K | 19 | 2.00% | 6 | 1.50% | 1 | 0.90% | 0 | 0.00% | 0 | 0.00% | 10 | 2.70% | 3 | 2.10% |
|  | Met | M | 31 | 3.30% | 15 | 3.60% | 3 | 2.80% | 2 | 3.50% | 1 | 2.30% | 11 | 3.00% | 4 | 2.80% |
|  | Phe | F | 30 | 3.20% | 11 | 2.70% | 1 | 0.90% | 1 | 1.80% | 0 | 0.00% | 13 | 3.50% | 5 | 3.50% |
|  | Pro | P | 75 | 8.10% | 38 | 9.20% | 12 | 11.20% | 5 | 8.80% | 7 | 16.30% | 26 | 7.00% | 11 | 7.70% |
|  | Ser | S | 109 | 11.70% | 59 | 14.30% | 12 | 11.20% | 2 | 3.50% | 9 | 20.90% | 36 | 9.80% | 14 | 9.90% |
|  | Thr | T | 52 | 5.60% | 21 | 5.10% | 4 | 3.70% | 0 | 0.00% | 3 | 7.00% | 25 | 6.80% | 6 | 4.20% |
|  | Trp | W | 12 | 1.30% | 3 | 0.70% | 0 | 0.00% | 0 | 0.00% | 0 | 0.00% | 7 | 1.90% | 2 | 1.40% |
|  | Tyr | Y | 20 | 2.20% | 6 | 1.50% | 1 | 0.90% | 0 | 0.00% | 1 | 2.30% | 11 | 3.00% | 3 | 2.10% |
|  | Val | V | 51 | 5.50% | 12 | 2.90% | 3 | 2.80% | 1 | 1.80% | 2 | 4.70% | 26 | 7.00% | 13 | 9.20% |
| MpARF2 | Ala | A | 59 | 6.70% | 19 | 5.30% | 0 | 0.00% | 0 | n/a | 1 | 3.00% | 34 | 8.60% | 2 | 1.70% |
|  | Arg | R | 57 | 6.50% | 23 | 6.40% | 4 | 7.50% | 0 | n/a | 1 | 3.00% | 27 | 6.80% | 6 | 5.10% |
|  | Asn | N | 32 | 3.60% | 16 | 4.50% | 4 | 7.50% | 0 | n/a | 4 | 12.10% | 14 | 3.50% | 4 | 3.40% |
|  | Asp | D | 52 | 5.90% | 15 | 4.20% | 2 | 3.80% | 0 | n/a | 1 | 3.00% | 22 | 5.60% | 10 | 8.50% |
|  | Cys | C | 12 | 1.40% | 3 | 0.80% | 1 | 1.90% | 0 | n/a | 0 | 0.00% | 7 | 1.80% | 1 | 0.80% |
|  | Gln | Q | 47 | 5.40% | 22 | 6.10% | 5 | 9.40% | 0 | n/a | 1 | 3.00% | 21 | 5.30% | 9 | 7.60% |
|  | Glu | E | 48 | 5.50% | 20 | 5.60% | 4 | 7.50% | 0 | n/a | 0 | 0.00% | 21 | 5.30% | 7 | 5.90% |
|  | Gly | G | 54 | 6.20% | 20 | 5.60% | 2 | 3.80% | 0 | n/a | 1 | 3.00% | 20 | 5.10% | 12 | 10.20% |
|  | His | H | 21 | 2.40% | 7 | 2.00% | 1 | 1.90% | 0 | n/a | 0 | 0.00% | 11 | 2.80% | 2 | 1.70% |
|  | Ile | I | 24 | 2.70% | 10 | 2.80% | 3 | 5.70% | 0 | n/a | 0 | 0.00% | 12 | 3.00% | 2 | 1.70% |
|  | Leu | L | 62 | 7.10% | 21 | 5.90% | 6 | 11.30% | 0 | n/a | 3 | 9.10% | 31 | 7.80% | 10 | 8.50% |
|  | Lys | K | 44 | 5.00% | 19 | 5.30% | 4 | 7.50% | 0 | n/a | 2 | 6.10% | 17 | 4.30% | 7 | 5.90% |
|  | Met | M | 18 | 2.10% | 8 | 2.20% | 0 | 0.00% | 0 | n/a | 0 | 0.00% | 8 | 2.00% | 2 | 1.70% |
|  | Phe | F | 26 | 3.00% | 8 | 2.20% | 1 | 1.90% | 0 | n/a | 0 | 0.00% | 14 | 3.50% | 5 | 4.20% |
|  | Pro | P | 72 | 8.20% | 33 | 9.20% | 3 | 5.70% | 0 | n/a | 4 | 12.10% | 35 | 8.90% | 4 | 3.40% |
|  | Ser | S | 120 | 13.70% | 62 | 17.30% | 6 | 11.30% | 0 | n/a | 11 | 33.30% | 42 | 10.60% | 14 | 11.90% |
|  | Thr | T | 45 | 5.10% | 22 | 6.10% | 3 | 5.70% | 0 | n/a | 1 | 3.00% | 19 | 4.80% | 6 | 5.10% |
|  | Trp | W | 14 | 1.60% | 5 | 1.40% | 0 | 0.00% | 0 | n/a | 0 | 0.00% | 7 | 1.80% | 2 | 1.70% |
|  | Tyr | Y | 16 | 1.80% | 4 | 1.10% | 2 | 3.80% | 0 | n/a | 1 | 3.00% | 8 | 2.00% | 4 | 3.40% |
|  | Val | V | 54 | 6.20% | 21 | 5.90% | 2 | 3.80% | 0 | n/a | 2 | 6.10% | 25 | 6.30% | 9 | 7.60% |

**Table S2.** *M. polymorpha* Mediator subunit orthologs.

|  | ID | Mp name | At ID | At Gene | Module |
| --- | --- | --- | --- | --- | --- |
| MED1 | n/a | n/a | n/a | n/a | Middle/Unclear |
| MED2/MED32 | Mp7g09990 | MpMED2 | AT1G11760 |  | Tail |
| MED3/MED27 | Mp4g22200 | MpMED3 | AT3G09180 |  | Tail |
| MED4 | Mp3g00880 | MpMED4 | AT5G02850 |  | Middle |
| MED5/MED24 | Mp1g09470 | MpMED5 | AT3G23590<br>AT2G48110 | REF4/RFR1 | Tail |
| MED6 | Mp8g03860 | MpMED6 | AT3G21350 | MHC9 | Head |
| MED7 | Mp6g01050 | MpMED7 | AT5G03220<br>AT5G03500 |  | Middle |
| MED8 | Mp1g02240 | MpMED8 | AT2G03070 |  | Head |
| MED9 | Mp3g14270 | MpMED9 | AT1G55080 |  | Middle |
| MED10 | Mp8g14670 | MpMED10 | AT5G41910<br>AT1G26665 |  | Middle |
| MED11 | Mp1g10270 | MpMED11 | AT3G01435 |  | Head |
| MED12 | MpUg00450<br>MpVg00710 | MpMED12U<br>MpMED12V | AT4G00450 | CCT | CDK |
| MED13 | Mp1g01940 | MpMED13 | AT1G55325 | MAB2/CFT2 | CDK |
| MED14 | Mp8g13300 | MpMED14 | AT3G04740 | SWP | Tail-Middle-Head |
| MED15 | Mp8g02180 | MpMED15 | AT1G15780 | NRB4/ATMED15a | Tail |
| MED16 | Mp4g19580 | MpMED16 | AT4G04920 | SFR6/GLH2 | Tail |
| MED17 | Mp5g24300 | MpMED17 | AT5G20170 |  | Head |
| MED18 | Mp2g10780 | MpMED18 | AT2G22370 |  | Head |
| MED19 | Mp5g04790 | MpMED19 | AT5G12230 |  | Head |
| MED20 | Mp6g06270 | MpMED20 | AT2G28230 |  | Head |
| MED21 | Mp8g06620 | MpMED21 | AT4G04780 |  | Middle |
| MED22 | Mp5g01850 | MpMED22 | AT1G16430<br>AT1G07950 |  | Head |
| MED23 | Mp1g05140 | MpMED23 | AT1G23230 |  | Tail |
| MED25 | Mp3g04170 | MpMED25 | AT1G25540 | PFT1 | Tail/Unclear |
| MED26 | Mp1g25310 | MpMED26 | AT3G10820<br>AT5G05140<br>AT5G09850 |  | Middle |
| MED28 | Mp4g00010 | MpMED28 | AT3G52860 |  | Head |
| MED30 | Mp3g02360 | MpMED30 | AT5G63480 |  | Head |
| MED31 | Mp4g01690 | MpMED31 | AT5G19910 |  | Middle |
| CDK8 | Mp2g10050 | MpCDK8 | AT5G63610 |  | CDK |
| CCNC | Mp1g05280 | MpCCNC | AT5G48640<br>AT5G48630 |  | CDK |

